# Metagenomic Profiles of Archaea and Bacteria within Thermal and Geochemical Gradients of the Guaymas Basin Deep Subsurface

**DOI:** 10.1101/2023.05.31.543050

**Authors:** David Geller-McGrath, Paraskevi Mara, Virginia Edgcomb, David Beaudoin, Yuki Morono, Andreas Teske

**Author notes:** These authors contributed equally (David Geller-McGrath, Paraskevi Mara).

## Abstract

While the temperature gradients of Earth’s crust and sediments are thought to delineate the downward extent and ultimate limits of the subsurface biosphere, the actual course of consecutively changing microbial communities and activities, from cool surficial sediments towards the deep, hot biosphere, remains to be charted. We used metagenomic and metatranscriptomic analyses of the hydrothermally heated, massive sediment layers of Guaymas Basin (Gulf of California, Mexico) to examine the environmental distribution and activity patterns of bacteria and archaea along thermal, geochemical and cell count gradients. Composition and distribution of MAGs, dominated by Chloroflexota and Thermoproteota, were shaped by biogeochemical parameters as long as temperatures remained moderate, but downcore increasing temperatures overrode other factors beyond ca. 45°C. Consistently, MAG genome size and diversity decreased with increasing temperature, indicating a conspicuous downcore winnowing of the subsurface biosphere. In contrast, specific archaeal MAGs within the Thermoproteota and Hadarchaeota increased in relative abundance and in recruitment of transcriptome reads towards deeper, hotter sediments, and mark the transition towards a distinct deep, hot biosphere.

## Introduction

The interplay between temperature stress and energy availability determines microbial survival in the subsurface biosphere, and delineates the extent and limits of life in the deep subsurface biosphere (Jørgensen and Hoehler 2013, Heuer et al. 2019). As microbial communities in cool, relatively shallow subsurface sediments transition into deeply buried and increasingly warm and finally hot sediments, it should be possible to track how subsurface bacteria and archaea react to these gradually harsher regimes on the levels of cellular activity and community change. While microbial abundance and diversity are expected to decline downcore (Starnawski et al. 2017, Kirkpatrick et al. 2019), it is also possible that particular subsurface-adapted microbial populations benefit from conditions that would eliminate others, and constitute a specialized deep, hot biosphere. Investigating downcore changes in microbial abundance, community composition and activity in well-characterized geochemical and thermal gradients requires a suitable field site where extensive physical, chemical and microbial gradients can be sampled in adequate resolution by sediment coring and drilling.

Guaymas Basin is a hydrothermally-active ocean spreading center in the central Gulf of California, covered by several hundred meters of sediment that host basaltic sill intrusions (Lizarralde et al. 2023) and strong geothermal heat flow (Neumann et al. 2023). Pyrolysis of buried organic carbon in these organic-rich sediments produces a complex milieu of petroleum hydrocarbons, including light hydrocarbons and methane, alkanes, and aromatic compounds, as well as carboxylic acids, and ammonia (Von Damm et al., 1985, Simoneit et al. 1995). These compounds are transported via hydrothermal fluids through Guaymas Basin’s thick sediments, supporting diverse and active microbial communities (Cruaud et al. 2017; Ramírez et al., 2021). Studies of surficial sediments that have been sampled by submersible show that sediment communities include filamentous nitrate-reducing, sulfur-oxidizing bacteria that form conspicuous mats on surfaces of hydrothermal sediments (Schutte et al. 2018), diverse methanogenic and methanotrophic archaea (Lever & Teske, 2015), thermophilic consortia of syntrophic sulfate-reducing bacteria and archaea that oxidize short-chain alkanes (Laso-Pérez et al., 2016; Krukenberg et al. 2016), and numerous anaerobic, often fermentative heterotrophs (Castro et al., 2021). Collectively, these communities not only perform chemosynthetic carbon fixation and heterotrophic organic matter remineralization, but they also assimilate fossil carbon into the benthic biosphere (Pearson et al., 2005).

Site to site variations in prokaryotic communities in sediments sampled by submersible are thought to be influenced by in situ temperatures and local geochemistry (Teske et al., 2021a). Few studies to date have explored the microbiology of deeper subsurface sediments in Guaymas Basin. Methanogens were enriched from sediments collected during Deep Sea Drilling Program Expedition 64 (Oremland et al., 1982), and bacterial and archaeal communities in piston cores were surveyed using 16S rRNA amplicon sequencing (Ramírez et al., 2020; Teske et al., 2019; Vigneron et al., 2014). Aside from these insights, the extent and characteristics of the deep biosphere in Guaymas Basin has remained largely unknown.

International Ocean Discovery Program (IODP) Expedition 385 drilled into Guaymas Basin at eight locations that differ in their degree of hydrothermal influence and heatflow (Neumann et al., 2023). Drilling sites followed broadly a northwest-to-southeast transect across the northern Guaymas axial valley (Figure 1). Two neighboring sites (U1545 and U1546) on the northwestern end of Guaymas Basin (Teske et al., 2021a, b) essentially differ by the presence of a massive, thermally equilibrated sill between 350 to 430 mbsf at Site 1546 (Lizarralde et al., 2023). Two drilling sites (U1547, U1548) targeted the hydrothermally active Ringvent area, approximately 28 km northwest of the spreading center (Teske et al., 2019), where a shallow, recently emplaced and hot sill creates steep thermal gradients and drives hydrothermal circulation (Teske et al., 2021c). Drilling Site U1549 (Teske, et al., 2021d) explores the periphery of an off-axis methane cold seep, Octopus Mound, located ∼9.5 km northwest of the northern axial graben (Teske et al., 2021e).

**Figure 1.**
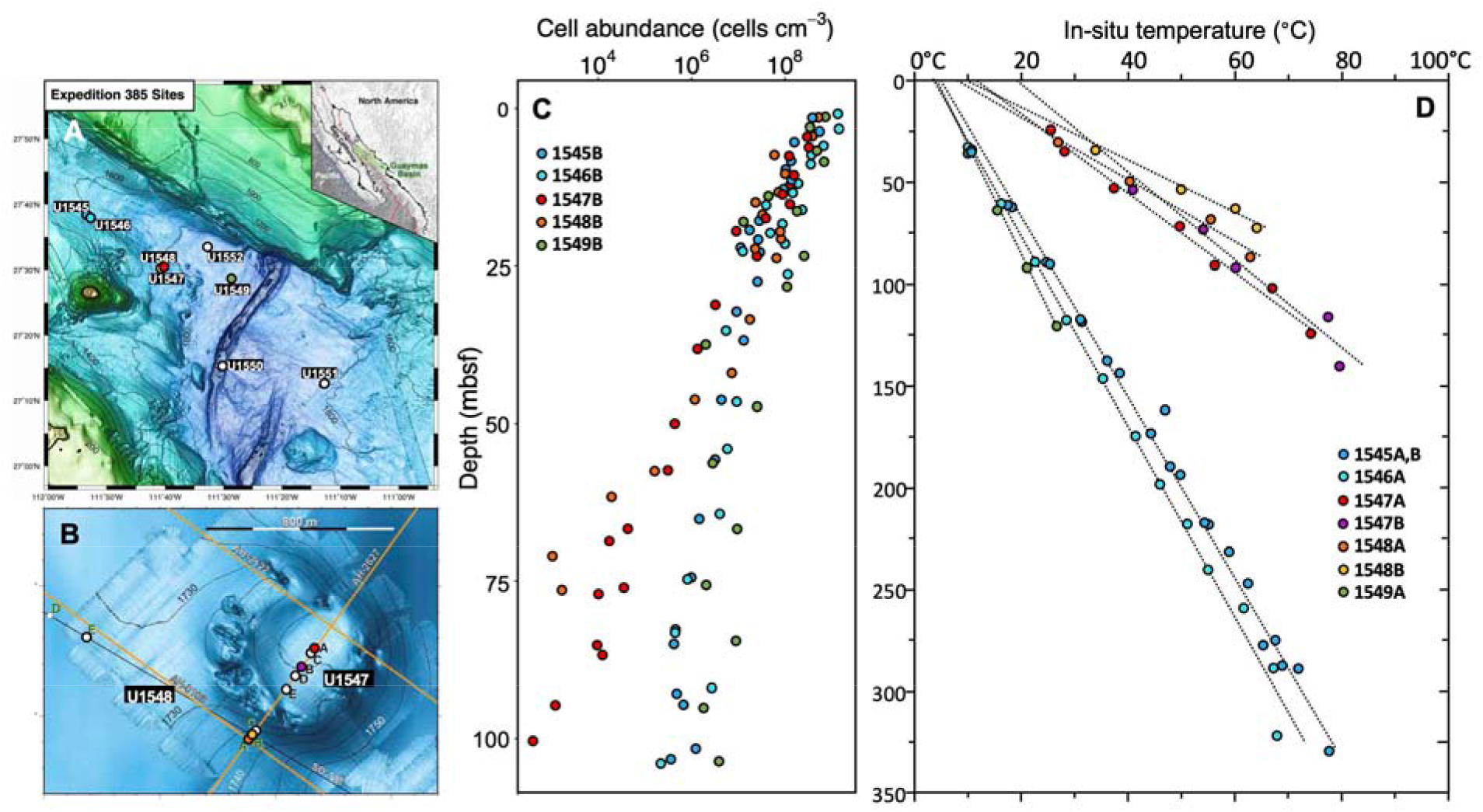
Locations, cell count profiles and temperature profiles for IODP Expedition 385 drilling sites. A) Guaymas Basin bathymetry with drill sites. B) Bathymetry of Ringvent with drill sites within and on the periphery of the Ringvent site. C) Cell counts for drill sites (U1545, U1546, U1547, U1548, and U1549) where metagenomic and metatranscriptomic samples were collected. D) Temperature profiles for drill sites where metagenomic and metatranscriptomic samples were collected. The lines indicated linear functions that were fitted to in-situ temperature measurements.

To assess the environmental distribution, genomic potential, subsurface adaptations and stress tolerances of microbes living in the deep biosphere of Guaymas Basin, we analyzed reconstructed metagenome-assembled genomes (MAGs) from depths ranging from 0.8 to 219.4 mbsf at these thermally and geochemically contrasting sites, and we provide mRNA evidence for the activity of selected bacterial and archaeal phyla.

## Results and Discussion

### Sampling sites and depths

Metagenomes were produced from sediment samples at drilling sites U1545B to U1549B that follow a northwest-to-southeast transect across the northwestern flanking region of Guaymas Basin (Figure 1A) and include an off-axis hydrothermal system, the Ringvent site (Figure 1B). The samples were selected to coordinate with depths used for separate ongoing analyses, and ranged from 1.7 m to 219 mbsf at Site U1545B, 0.8-16.3 mbsf at U1546B, 2.1-75.7 mbsf at U1547B, 9.1-69.4 mbsf at U1548B, and 16.5 mbsf at U1549B (Figure 1; Table 1). For all samples, a wide range of geochemical parameters was analyzed shipboard (Supplementary Table 1). The sites represent distinctly different thermal gradients and cell densities; generally, sites with steeper downcore temperature gradients are characterized by more rapidly decreasing cell counts (Figure 1C,D). U1545B is the reference site for IODP Expedition 385 because of the absence of seepage, hydrothermal influence, and massive sill intrusions (Teske et al., 2021a). Here, metagenome libraries extended down to 219.4 mbsf, at in-situ temperatures of 54°C. Cell count trends for site U1545, U1546 and U1549 were similar, and showed a decrease over three orders of magnitude within 100 meters (Figure 1C). At the hot Ringvent sites U1547B and U1548B (Neumann et al. 2023), comparable temperatures of 50-55°C were already reached near 70 mbsf (Figure 1D), and cell counts decreased by at four to five orders of magnitude within this depth range (Figure 1C). To describe temperature-related trends in MAG recovery and diversity, we categorized our samples into three groups according to temperature, cool (2-20°C), warm (20-45°C) and hot (45-60°C).

### Subsurface Biogeochemical zonation

Most metagenome samples are from sediments within the sulfate-reducing zone where sulfate is still available at near-seawater concentrations (∼28 mM) or becomes gradually depleted with depth. At those same sediments hydrogen sulfide concentrations are gradually increasing towards multiple millimolar values. Metagenome samples from site U1545B also include depths spanning the sulfate-methane transition zone (SMTZ) at ∼ 64 mbsf; at the SMTZ sulfate concentrations drop from 21.1 mM to 0.7 mM, sulfide reaches peak concentrations of 8.9 mM, and methane concentrations increase from picomolar to 1.5 mM (Table 1). High methane concentrations persist also in deeper samples from U1545B, and decrease only in the very deepest samples (> 200 msbf). Samples from Ringvent sites (U1547B and U1548B) show gradual, but incomplete downcore sulfate depletion (from 27.9 to 18.8 mM) combined with hydrogen sulfide accumulation (max. 7.1 mM at 75.7 mbsf at U1547B). Ammonia concentrations increase from < 1 mM to 3-5 mM downcore at most sites, and reach 9-25 mM below the SMTZ in U1545B. Dissolved inorganic carbon (DIC) and alkalinity concentrations are generally highest at site U1545B where they peak in the SMTZ (∼28 and 60 mM, respectively). Ammonia, DIC and alkalinity remain elevated in the deeper samples of site U1545B, presumably due to cumulative bioremineralization over time. In contrast, the Ringvent samples generally have lower ammonia, alkalinity and DIC porewater concentrations, suggesting reduced bioremineralization of organic matter at these sites. Dissolved organic carbon (DOC) remained ∼10 to 20 mM in most samples but increased towards 70 mM in the sulfate-methane transition zone of U1545B and remained between 20 and 50 mM in the deeper sediments of U1545B. This suggests DOC enrichment and decreased heterotrophic DOC consumption in deep methanogenic sediments of U1545B where energy-rich electron acceptors for heterotrophic carbon remineralization are not available. Total nitrogen and total organic carbon decrease with depth at all sites (Table 1). Total petroleum hydrocarbon content in surficial sediments ranges from 150 to 250 µg/g sediment and decreases slightly downcore, before increasing considerably (sometimes into the mg/g range) in hot sediments near deep sill intrusions (Supplementary Figure 1).

### MAG diversity, distribution, and evidence of activity

A total of 142 metagenome-assembled genomes (MAGs) were recovered from a co-assembly of all metagenomic samples. MAGs that matched those from negative controls were excluded from further analysis (Supplementary Table 2). For downstream analysis, we retained 89 MAGs that had at least ≥ 50% bin completeness and ≤ 10% bin contamination (Supplementary Table 3). Genome completeness ranged from ∼50 to 97%. (Supplementary Figure 2; Bowers et al. 2017).

Taxonomic assignment showed that the 89 MAGs were assigned to 6 archaeal and 13 bacterial phyla (Figure 2, Supplementary Figure 3) that overlapped with lineages documented previously in 16S rRNA gene amplicon sequencing of shallow subsurface sediments (Ramírez et al. 2020, Teske et al. 2019), and in metagenomic surveys of shallow hydrothermal sediments of Guaymas Basin (Dombrowski et al. 2018). In parallel to downcore decreasing cell numbers (Figure 1), MAG diversity decreased downcore at all sites as temperatures increased (Figure 2). In samples from cool (3-20°C) sediments from all sites, reads mapped to diverse bacterial and archaeal phyla, including the bacterial phyla Chloroflexota, Acidobacteriota, Desulfobacterota, WOR-3, Aerophobota, and Bipolaricaulota, and the archaeal phyla Thermoproteota, Thermoplasmatota, and Aenigmatarchaeota (Figure 2). In samples with warm temperatures (20-45°C), reads were predominantly assigned to bacterial phyla Chloroflexota (Order G1F9), Acidobacteriota, WOR-3 (UBA3073), Aerophobota and Bipolaricaulota, and to archaeal phyla Thermoproteota, Hadarchaeota, and Aenigmatarchaeota. At hot temperatures (45-60°C), bacterial reads mapped primarily to a single Chloroflexota MAG (class Dehalococcoidia), a single WOR-3 MAG and two Aerophobota MAGs (class Aerophobia). In contrast, several Archaeal MAGs show a marked preference for hot sediments, and mapped to the Thermoproteota (class Bathyarchaeia), and Hadarchaeota (class Hadarchaeia). Our recovered MAGs reflected metabolisms predicted for the deep biosphere including fermentation, sulfate reduction, alkane degradation, and carbon fixation. In addition, the potential for iron reduction using extracellular electron transfer mechanisms (e.g., *mtrA, mtoA,* and *eetB* genes; Garber et al. 2020; Chatterjee et al. 2021) was found in 86/89 MAGs from all recovered phyla (Supplementary Tables 4-5, and Supplementary Text). Aside from genes associated with metabolic processes, we note the presence of CRISPR/Cas and CRISPR *Csm* genes involved in viral defense-associated DNA editing, e.g. in Aerophobota MAGs distributed across thermal regimes.

**Figure 2.**
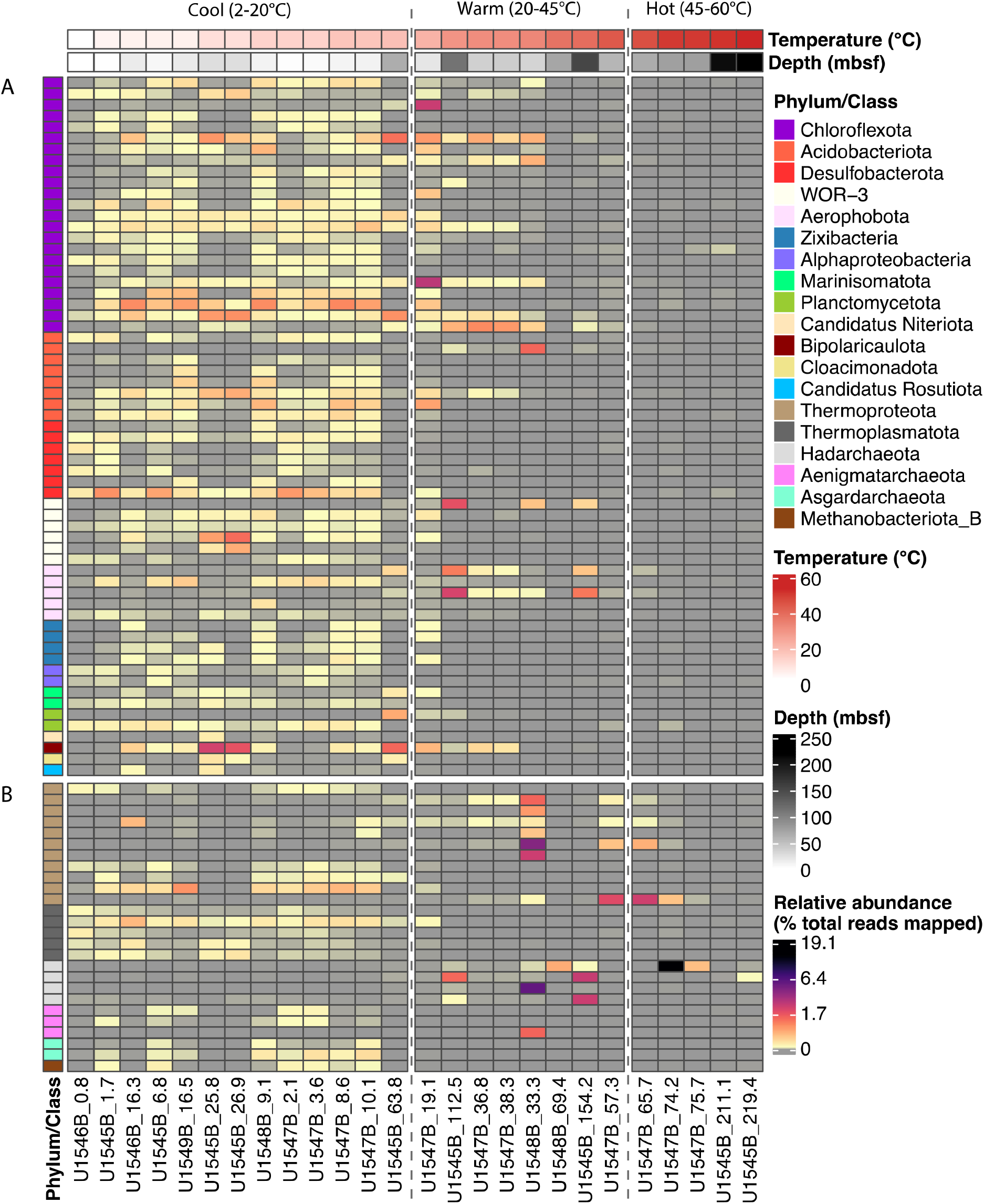
Heatmap of MAG relative abundance. Each column shows the percentage of total pre-processed reads from a metagenomic sample that mapped to all 89 MAGs across all samples, in order of increasing temperature from left to right. Panel **A** denotes bacterial MAGs and panel **B** denotes archaeal MAGs. Temperature regimes (Cool, Warm, and Hot) are separated by vertical dashed lines. Each row represents the abundance profile of an individual MAG across all samples. MAGs are color-coded by phylogenetic affiliation. Samples are color coded according to increasing temperature from left to right. The x-axis sample labels indicate site and depths in mbsf. Abundance values are the percent of total metagenomic reads per sample that mapped to each MAG.

To determine any intra-phylum differences in metabolic activity, we mapped metatranscriptome reads to our recovered MAGs, for samples collected at the same sites (Mara et al., submitted). Since the metagenome and metatranscriptome of the Guaymas Basin subsurface remain incompletely covered by sequence data, the absence of transcript read mapping to particular MAGs cannot be taken as evidence of microbial inactivity. Microbial activity of the deep biosphere is certainly constrained but not eliminated by substrate and energy limitation (Hoehler & Jørgensen, 2013). To avoid these ambiguities that are inherent in negative transcript mapping results, we focus further discussion on positive transcript mapping results that support the activity of specific MAGs in the subsurface. Actively transcribed genes are present for MAGs within all phyla discussed here, albeit at variable levels; some MAGs within individual phyla show no or much lower apparent activity than others (Figure 3). Transcript mapping revealed widely distributed metabolic capabilities that are anticipated in the deep biosphere, including chemoautotrophy, sulfate reduction, fermentation, and hydrogen utilization (Supplementary text). Bacterial transcripts were mapped predominantly within the phylum Chloroflexota. In our hot sediment samples, most transcripts were mapped to archaeal MAGs, within the phyla Thermoproteota, Hadarchaeota, and Aenigmatarchaeota (Figure 3).

**Figure 3.**
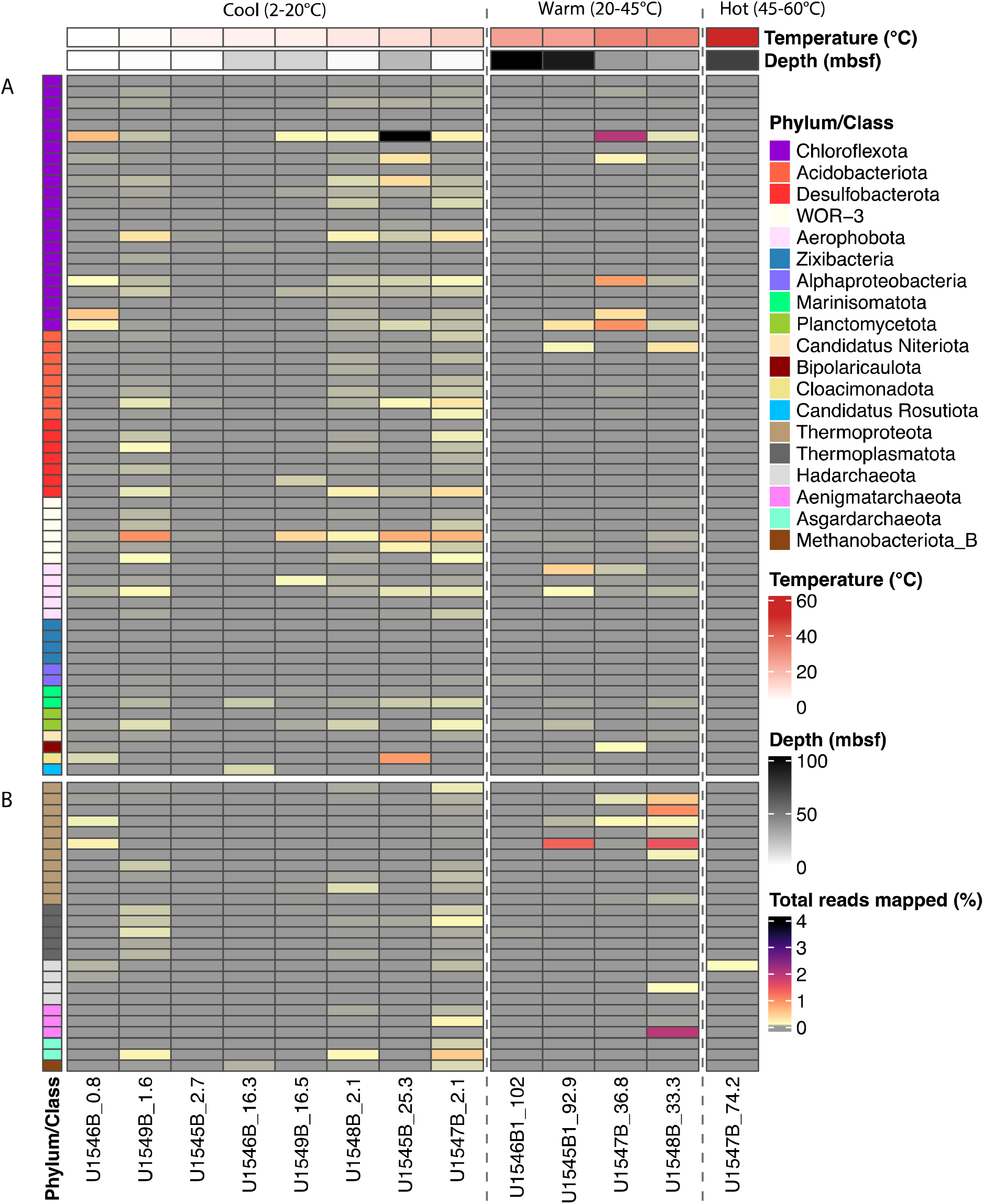
Heatmap of MAG Metatranscriptomic read recruitment. Each column shows the percentage of total pre-processed reads from metatranscriptomic samples that mapped to all 89 MAGs across all samples, in order of increasing temperature from left to right. Panel **A** denotes bacterial MAGs and panel **B** denotes archaeal MAGs. Temperature regimes (Cool, Warm, and Hot) are separated by vertical dashed lines. Each row represents the abundance profile of an individual MAG across all samples. MAGs are color-coded by phylogenetic affiliation. Samples are color coded according to increasing temperature from left to right. The x-axis sample labels indicate site and depths in mbsf. Abundance values are the percent of total metatranscriptomic reads per sample that mapped to each MAG.

### The influence of environmental factors on MAG composition

The relationship between environmental parameters and the taxonomic composition of MAGs from cool (2-20°C), warm (20-45°C), and hot temperatures (45-60°C) was investigated using non-metric multidimensional scaling (nMDS). Bacterial and archaeal communities clustered by temperature (Figure 4; Supplementary Figure 4), comparable to previous analyses of temperature-dependent microbial community composition in surficial sediments in Guaymas Basin (Teske et al., 2021d). Fitting a suite of environmental variables to our nMDS analyses revealed carbon monoxide (CO), total organic carbon content (TOC), total sulfide (H_2_S), total nitrogen content (TN), pH, salinity, phosphate concentration (PO ^3-^) and temperature to have a significant effect (p < 0.05) on taxonomic composition of MAGs in samples. MAG composition within the cool temperature range was influenced most strongly by concentrations of CO, TOC and TN. Within the warm temperature range, pH, salinity, TOC and H_2_S affected MAG composition. For MAGs recovered in the hot temperature range, temperature in itself was the strongest influencing factor. The influence of TOC and TN on MAG diversity in cool samples may reflect increased availability of labile sources of dissolved and particulate organic carbon in near-surface sediments, in contrast to more recalcitrant substrates in deeper and warmer subsurface sediments. Similarly, increasing H_2_S concentrations in downcore samples create an extremely reductive environment that may select for specific phyla (e.g., Hadarchaeota) or specific taxa within other detected phyla. For MAGs from hot samples, carbon substrates or other chemical parameters becomes secondary to the impact of temperature itself.

**Figure 4.**
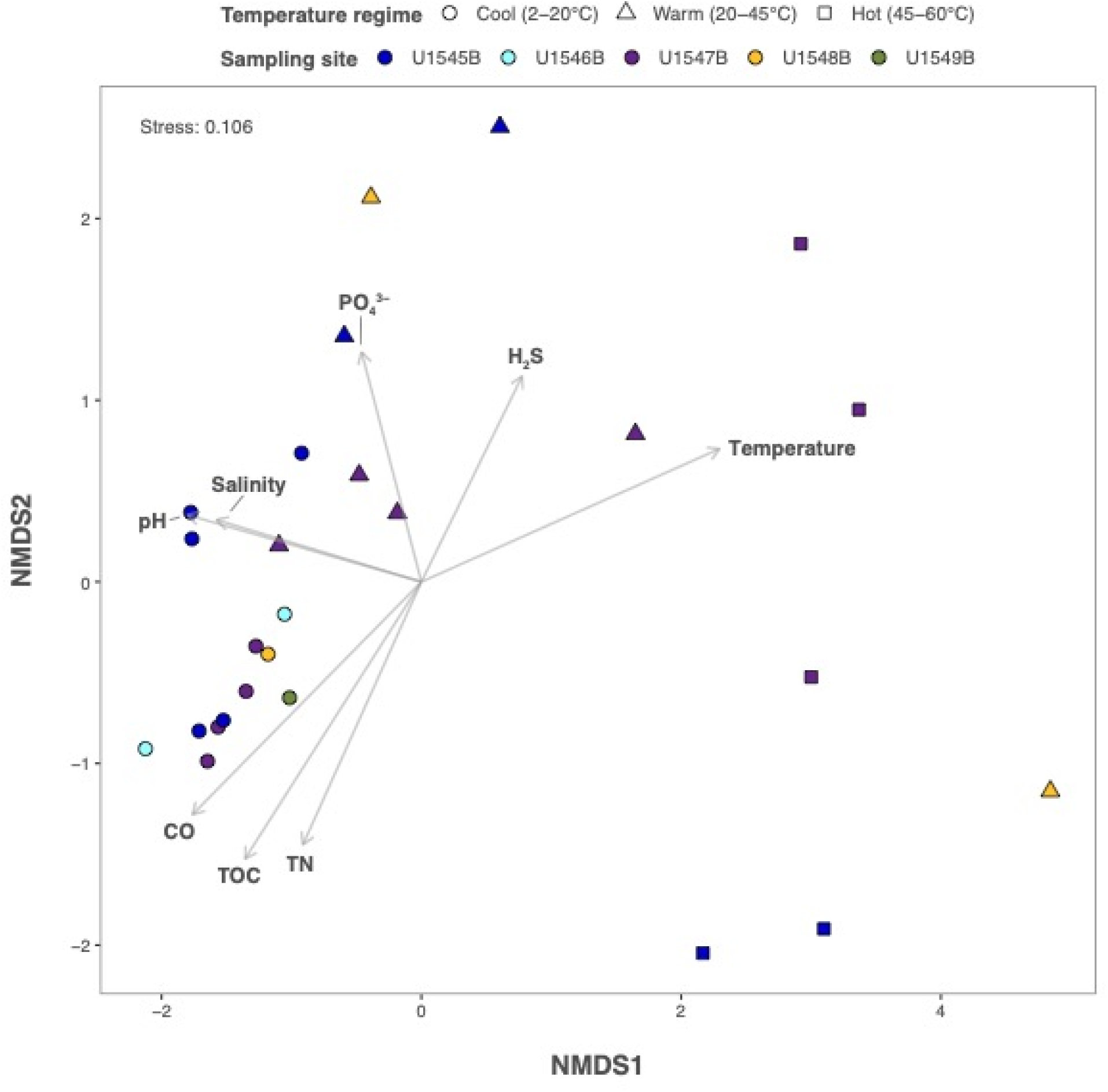
Non-metric multidimensional scaling (nMDS) ordination plot of metagenomes reconstructed from sites U1545B, U1546B, U1547B, U1548B, U1549B and environmental parameters with a significant effect on microbial community composition. The nMDS plot depicts the separation of Guaymas Basin metagenomic samples drilled from depths spanning 0.8-219 mbsf and temperatures ranges for cold (0-20°C), warm (20-45°C) and hot (45-60°C) sediments, and statistically significant (p < 0.05) environmental parameters; plot stress: 0.106. The directions of the arrows indicate a positive or negative correlation among the environmental parameters with the ordination axes. The length of the arrow reflects the strength of correlation between the environmental parameter and the microbial community composition of metagenomic samples, with longer lines indicating stronger correlations.

### Metagenomic subsurface adaptations

In addition to genes for core metabolic processes (e.g., glycolysis, biosynthesis of nucleotides and amino acids), Guaymas Basin MAGs contain widespread genomic features extend across multiple bacterial and archaeal phyla that appear to have adaptive value for survival in the Guaymas Basin subsurface. These include a variety of transporters and efflux pumps associated with microbial defense, biosynthetic gene clusters involved in synthesis of diverse secondary metabolites, and two-component systems (TCSs) that can induce metabolic shifts.

TCSs are used extensively by bacteria and some archaea to respond and adapt to environmental changes (Beier & Gross, 2006); archaea acquire TCS genes through horizontal gene transfer from bacteria (Schaller et al., 2011). In Guaymas Basin, TCS genes occur in the majority of bacterial MAGs but not in archaeal MAGs (Supplementary Tables 4-5), and they may help cells to adapt to long term burial. For example, the KinABCDE-Spo0FA system is present in almost all our bacterial MAGs and plays a role in sporulation by shifting cellular metabolism from active growth to dormancy/sporulation (Quisel et al., 2001). Likewise, the RegB/RegA redox-signaling mechanism involved in carbon fixation, hydrogen oxidation and anaerobic respiration (Elsen et al., 2004) is present in the majority of the bacterial MAGs.

Bacterial and archaeal MAGs encoded transporters and efflux pumps associated with microbial defense (e.g., multidrug resistance pumps), and biosynthetic gene clusters involved in the biosynthesis of diverse secondary metabolites (Supplementary Table 6). Biosynthetic gene clusters associated with archaea were primarily annotated as polyketide synthases, ribosomally synthesized and post-translationally modified peptides. Additionally, KEGG modules identified archaeal genes encoding terpene synthesis (e.g., geranylgeranyl diphosphate synthase; Supplementary Table 5) that have diverse functions (e.g., pigments, antimicrobial agents and in plants as thermoprotectants; Yang et al. 2012).

Genes involved in chemotaxis (*cheA*/*B/R/W/Y*) and motility (*flgB/C/E/G/H/I* and *fliE/F/G/*) were present in bacterial and archaeal MAGs. These findings suggest widespread potential for cell-cell interaction, cell movement and competition for resources in the Guaymas Basin subsurface microbial community. The expression of genes associated with secondary metabolite biosynthesis and cell motility in marine subsurface MAGs has been reported previously for Peru Margin and Canterbury Basin sediments at depths up to 345 mbsf (Pachiadaki et al., 2016). Other metagenomic studies have argued that cell motility genes are depleted in the marine subsurface (Biddle et al., 2008), therefore determining the functions of these genes requires further investigation.

### Characteristics and distribution of dominant bacterial and archaeal groups

We examined MAGs affiliated with dominant phylum-level lineages (Chloroflexota, Thermoproteota, Hadarchaeota) with distinct mesophilic and thermophilic preferences, and examined their order-level lineages that appear at specific depth and temperature ranges. An extended overview on MAGs from these and other bacterial and archaeal phyla is provided in the supplementary materials.

### Chloroflexota

Chloroflexota are the dominant bacterial phylum in the Guaymas subsurface MAGs and account for 23 out of 89 unique MAGs, comprise 12 order-level lineages, and account for a significant fraction of recruited metagenomic reads per sample (up to 8.3%) (Figure 2, Supplementary Figure 5). The metabolic potential of MAGs recovered for this diverse phylum is detailed in the Supplementary text. This phylum thrives in the deep biosphere and includes metabolically diverse heterotrophs within thermophilic and dehalogenating lineages. Chloroflexota are abundant in the hadal ocean (Liu et al., 2022a), in marine subsurface sediments (Vuillemin et al., 2020) and hydrothermal settings (Fullerton & Moyer, 2016, Dombrowski et al., 2018; Reysenbach et al., 2020). At site U1545B, almost all Chloroflexota MAGs were detected within cool shallow samples (1.7-6.8 mbsf and ∼6°C). Order-level lineages within the Chloroflexota changed from members of the widely distributed Groundwater InFlow clone 9 (GIF9) group (Frouin et al., 2018; Hug et al., 2013) at 25.8 mbsf (11°C) and at the SMTZ (63.8 mbsf, 19°C), to MAGs affiliated with the Dehalococcoidales lineage at 112.5 and 154.2 mbsf (30 and 40°C, respectively). At Ringvent site 1547B, Chloroflexota MAGs were abundant in samples between 2.1-10.1 mbsf (∼14-17°C), declined at 19.1 mbsf (22°C), and became undetectable at 57.3 mbsf (45°C) (Supplementary Figure 5). Although diverse thermophilic Chloroflexota lineages have been enriched and isolated from hot springs (Dodsworth et al., 2014; Palmer et al., 2023), the Guaymas Basin subsurface Chloroflexota appear to prefer cool or moderately warm habitats within the upper 60 mbsf, and avoid temperatures above 30-40°C.

### Thermoproteota

Archaeal MAGs were dominated by the Thermoproteota; all 11 MAGs within this phylum belonged to the class Bathyarchaeia, one of the four archaeal lineages – in addition, Thaumarchaeota, Aigarchaeota, and hyperthermophilic Korarchaeota - that constitute this phylum (Oren & Garrity, 2021). Bathyarchaeial MAGs were detected at all examined sites but primarily at the hot Ringvent sites U1547B and U1548B (Figure 2 and Supplementary Figure 5). Five MAGs assigned to the orders of TCS64 and 40CM−2−53−6 were recovered primarily between 0.8-15 mbsf sediments at sites U1545B and U1547B with cool temperatures ranging from 2.8-17.4°C. These bathyarchaeial orders have been reported also from deep sea brine pool samples (Zhang et al., 2016) and from soil samples (Butterfield et al., 2016). The order-level lineage B26-1, previously found in Guaymas Basin hydrothermal sediments collected via *Alvin* pushcores (Dombrowski et al., 2018; He et al., 2016), included three MAGs from warmer sediments (19-39.5°C) below 63.8 mbsf at U1545B, and warm to hot sediments (22-47°C) between 19.1 and 65.68 mbsf at Ringvent site U1547B. The order-level lineage RBG-16-48-13, recovered previously from terrestrial subsurface drill cores (Anantharaman et al., 2016), was represented by a MAG detected at site U1548 at 20-45°C (Suppl. Fig. 3). Two bathyarchaeial MAGs could not be classified at the order level, but one of these MAGs was abundant at temperatures between 39.5-51°C at U1547B (Supplementary Figure 5). The detection of bathyarchaeial MAGs over a wide temperature spectrum, and the link of bathyarchaeial orders to specific temperature regimes, is consistent with distinct thermal preferences among different lineages of Bathyarchaeia (Qi et al., 2021). Bathyarchaeia widely occur in marine subsurface sediments (Lloyd et al., 2013), including hydrothermal sediments of Guaymas Basin (He et al., 2016). Their ubiquitous presence in anaerobic and hydrothermal sediments can be attributed to their capacity to metabolize multiple organic substrates, e.g., polysaccharides, urea, detrital proteins, aromatics compounds such as benzoate and lignin, and acetate (Lazar et al., 2016; Lloyd et al., 2013; Seyler et al., 2014, Yu et al. 2018). The capacity of Bathyarchaeia to degrade substituted aromatic compounds might contribute to their persistence in the hydrocarbon-rich Guaymas Basin subsurface. Based on MAG gene content, Bathyarchaiea can potentially utilize formaldehyde and shuttle it into carbon fixation via the Wood-Ljungdahl pathway (see Supplementary Text). Lineage-specific thermophilic adaptations among the Bathyarchaeia include reverse DNA gyrase that facilitates DNA supercoiling under extreme temperatures (Feng et al., 2019).

### Hadarchaeota

Hadarchaeota are found widely in subsurface sediments where they survive by a combination of heterotrophic traits (fermentation of sugars) with autotrophic energy generation, specifically the oxidation of carbon monoxide and hydrogen (Baker et al., 2016). Hadarchaeota were previously recovered from surficial hydrothermal sediments in Guaymas Basin (Dombrowski et al., 2018). The majority of metagenomic reads from depths below 60 mbsf at sites U1545B and U1548B were recruited to 4 MAGs affiliated with Hadarchaeota (order Hadarchaeales) (Figure 2). These Hadarchaeota MAGs did not recruit any reads from cool samples but were detected only in warm and hot samples, indicating a preference for elevated temperatures (Supplementary Table 1). Thus, the Hadarchaeota, originally detected in hot aquifers of deep terrestrial mines (Takai et al., 2001), emerge as the best archaeal candidate with a consistent preference for elevated temperatures in deep sediments of the Guaymas Basin subsurface.

One Hadarchaeota MAG, Guaymas_P_008 (92% completeness, 4% contamination), appears particularly abundant at 74.2 msbf at the hot Ringvent site U1547B where *in situ* temperature was 51°C, recruiting ∼19% of reads from that sample (Figure 2 and Figure 3). This MAG contained carbohydrate hydrolysis genes (α-RHA, β-galactosidase) and nucleoside uptake and degradation marker genes (nucleoside transporters, purine nucleosidases) that suggest they utilize nucleosides for purine/pyrimidine synthesis. This MAG also contained carbon monoxide oxidation genes (*coxM*, *coxS*) that were absent in the other Hadarchaeota MAGs that primarily encoded genes for fermentation (*porA*, *ack*, *acdA*) and aromatics degradation (*ubiX*) (Supplementary Tables 5-6). The ability to utilize a wider range of carbohydrates may contribute to its success at higher temperatures, as has been reported previously for thermally-adapted Bathyarchaeia genomes (Qi et al. 2021). Considering the potential for hydrocarbon utilization in the Hadarachaeota and other phyla (Supplementary Text), microbial utilization might contribute to reduced hydrocarbon concentrations at intermediate sediment depths and temperatures (Supplementary Figure 1). One of our Hadaracheota MAGs (P_034) contained *mcrC* and *mcrG* homologs that regulate the expression and assembly of the alkyl/methyl coenzyme M reductase operon (Shao et al. 2022), the essential methane and alkane-activating genes in archaeal methanogens, methane oxidizers and short-chain alkane oxidizers (Wang et al., 2021). Aside from metabolic genes, we note the presence of KaiC histidine in Hadarchaeota, a circadian clock protein that regulates cell division and allows prokaryotes to adapt to changes in environmental conditions (Jabbur and Johnson, 2022) and the gene for programmed cell death protein 5, that is linked to anti-virus defense and triggers dormancy under hostile conditions (Koonin et al., 2017).

### Genome sizes of subsurface MAGs

Comparisons of estimated genome sizes for all MAGs that recruited at least 0.1% of metagenomic reads from cool, warm, and hot sediments revealed a difference in average genome size. The most abundant genomes in cool sediments were on average significantly (two-sided Welch’s *t*-test, adjusted p < 0.05) larger (∼32%) than those recovered from hot sediments (Figure 5A,B). The estimated genome size of MAGs recovered from our shallow (2-15 mbsf) samples was also ∼22% larger on average than those detected in deeper (> 60 mbsf) and warmer (>30-40°C) sediments. Linear regression analysis demonstrated a general reduction in average genome size in our samples as both temperature and depth increased (Figure 5C, D). Elevated *in situ* temperatures are thought to select for smaller genome sizes via genome streamlining (Sabath et al. 2013), for example increased gene loss after duplication; the effects of genomic streamlining are pervasive and result in the elimination of hundreds of genes all over the genome (Simonsen 2022). Reduced genome size lowers the metabolic cost required for microbial DNA replication, as suggested for hadal microorganisms in the Challenger Deep at Mariana Trench (Zhou et al. 2022). Microbes with smaller genomes would gain a relative survival advantage and gradually dominate the microbial community in the subsurface, as metabolically more demanding microbial community members with large genomes die off. Such a mechanism would be accompanied by microbial community change; it would contribute to the selection of subsurface-adapted microbial communities that has been documented already within the top few meters below seafloor (Starnawski et al. 2017), and it would explain the small size of microbial cells in deep subsurface sediments, near 0.5 micrometer (Biddle et al. 2006).

**Figure 5.**
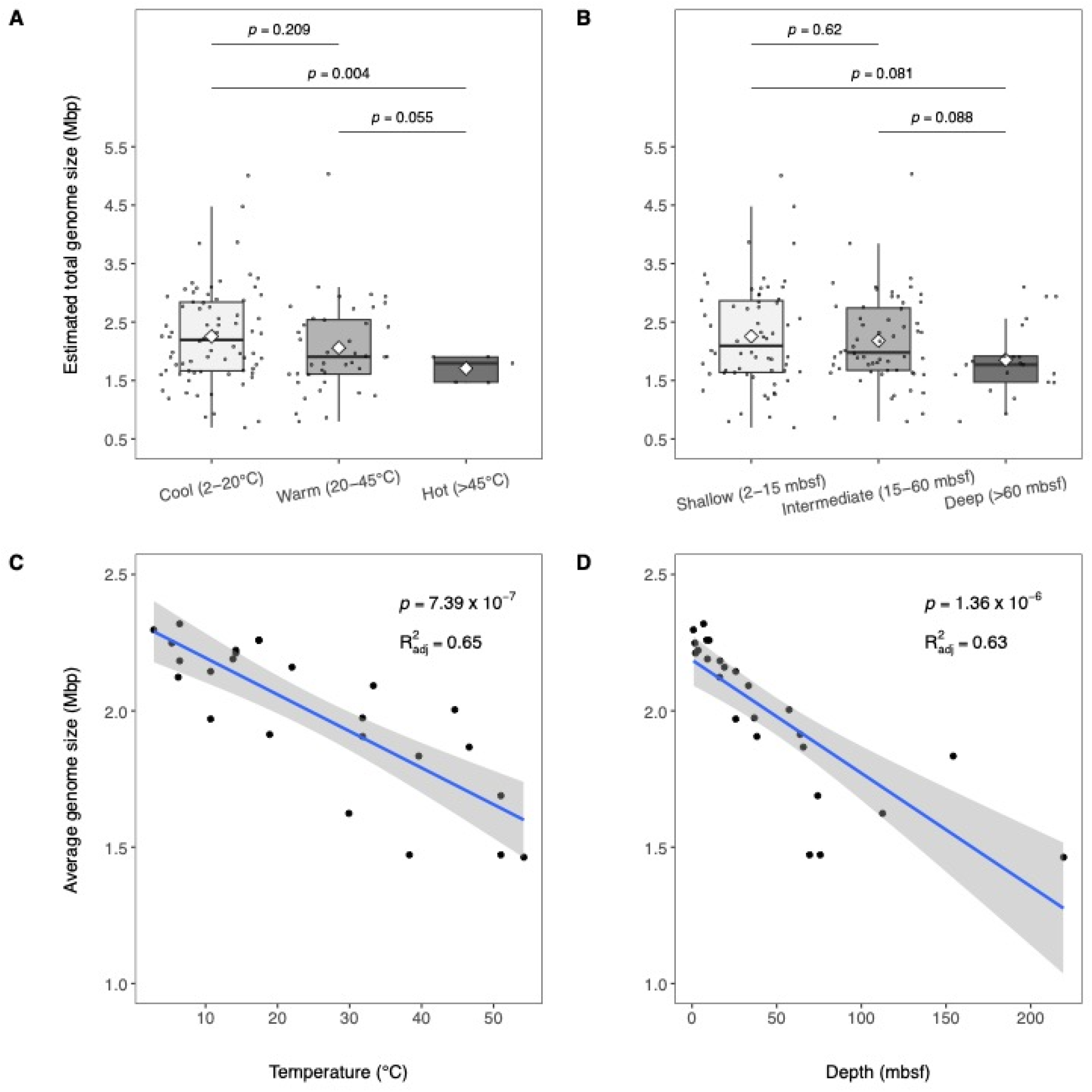
Estimated and average genome size vs. temperature and depth. Boxplots show estimated genome size of MAGs that recruited at least 0.1% of metagenomic reads from samples collected in cool (2-20°C), warm (20-45°C), and hot sediments (45-60°C) in panel **A,** and at shallow (2-15 mbsf), intermediate (15-60 mbsf), and deep (>60 mbsf) depths in panel **B**. The Median is shown the middle horizontal lines, the mean as the white diamonds, and interquartile ranges are shown as boxes (whiskers extend to 1.5 times the interquartile range). Each data point is overlaid on the boxplots and values at the top denote adjusted p-values from two-sided Welch’s *t*-tests comparing estimated genome sizes by temperature (panel **A**) and depth (panel **B**) regimes. Panels **C** and **D** display the relationship between average estimated genome size in each metagenomic sample plotted against temperature (**C**) and depth (**D**) using linear regression. The blue lines in panels **C** and **D** denote the regression lines, with the 95% confidence intervals indicated by the grey bands. The values at the top of panels **C** and **D** denote the p-value and adjusted R-squared value of the fit.

### Subsurface MAG recovery limits

The environmental stresses that increasingly exclude microbial lineages and reduce genome size are finally reflected in decreased recovery of unique MAGs in warmer and deeper samples from all sites (Figure 6A and B, respectively). Plotted against depth, MAG recovery declines more quickly for the two hotter Ringvent sites U1547 and U1548 than for the cooler sites U1545, U1546 and U1549 (Figure 6B). When plotted against temperature, declining MAG recovery for the hot Ringvent sites and the cooler sites converge towards a shared minimum between ca. 50 and 55°C (Figure 6A). These comparisons show that the decline of MAG recovery with depth is locally modified, behaves differently at different sites, and does not follow a uniform depth-related decay rate. In contrast, the influence of increasing temperature is pervasive, reduces microbial diversity at all sites, and occludes the emergence of MAGs representing new microbial lineages beyond approximately 50-55°C.

**Figure 6.**
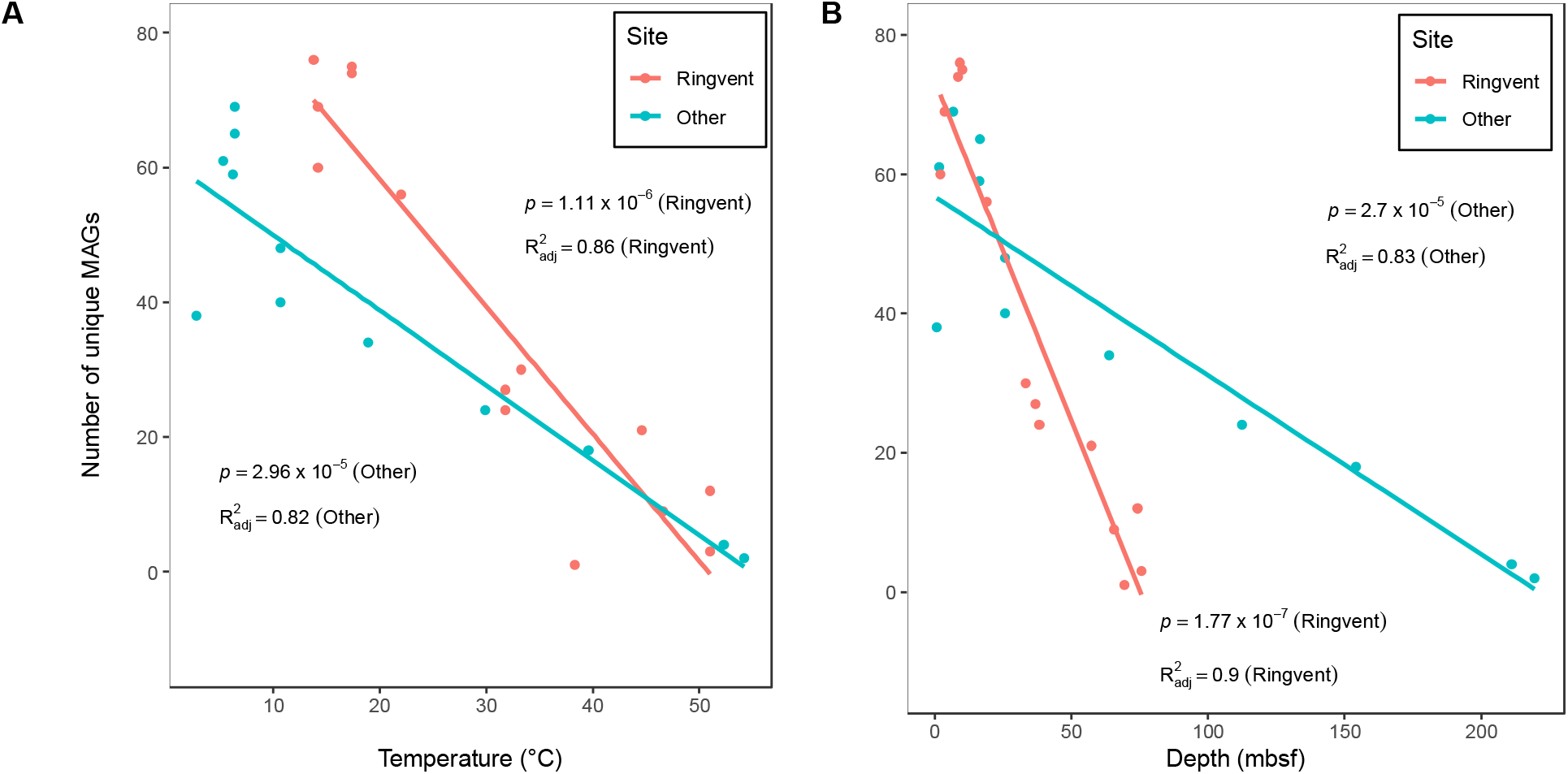
MAG recovery at sampled sites vs. temperature and depth. Panels **A** and **B** display the relationship between the number of unique MAGs detected in samples from sites U1547 and U1548 (in red) and sites U1545, U1546, and U1549 (in green) plotted against temperature (**A**) and depth (**B**) using linear regression. The solid lines in panels **A** and **B** denote the regression lines, with the 95% confidence intervals indicated by the grey bands. For both panels **A** and **B** the p-values and adjusted R-squared values of the fits of each regression line are shown.

## Conclusions

While improved DNA and RNA recovery could potentially overcome declining downcore cell density, and extend the recovery of new bacterial and archaeal MAGs towards deeper and hotter sediments, the observed trend towards increasingly limited microbial diversity in the subsurface stands in marked contrast to the numerous bacterial and archaeal lineages that thrive in surficial hydrothermal sediments of Guaymas Basin, where fluidized sediments are permeated by pulsating, extremely hot (> 80°C) and highly reducing fluids (Dombrowski et al. 2018). We ascribe the difference to contrasting energy supply, and suggest that relatively moderate temperatures in IODP boreholes have a disproportionally greater impact on the energy-limited microbial deep biosphere than hydrothermal temperatures have on energy-replete hydrothermal communities in surficial sediments. The latter conditions select for diverse thermophilic and hyperthermophilic, frequently chemolithoautotrophic bacterial and archaeal groups (Anderson et al. 2015). We suggest that this difference ultimately results in distinct microbial communities in subsurface sediments and surficial hydrothermal sites of Guaymas Basin (Lagostina et al. 2021), and accounts for the significant overlap of Guaymas Basin subsurface bacteria and archaea – predominantly Chloroflexota, Thermoproteota, Acidobacteriota, and Desulfobacterota - with the largely heterotrophic and mesophilic microbiota of non-hydrothermal benthic sediments (Baker et al. 2021, Parkes et al., 2014). Yet, we note that specific archaea, in particular the Hadarchaeota, show a preference for deep, hot sediments of Guaymas Basin; these archaea extend further into the sediment column, beyond the current limit of MAG detection (Paraskevi et al., in review), and appear to represent deep subsurface thermophiles that are sustained by substrates and energy sources of deep, hot sediments. Observations of microbial cells and activity in extremely deep, hot subsurface environments (Inagaki et al. 2015, Heuer et al. 2020) could indicate such thermophile communities that have adapted to deep subsurface conditions.

## Methods

### Sample collection

Sediment cores were collected during IODP Expedition 385 using the drilling vessel *JOIDES Resolution*. Holes at each site were first advanced using advanced piston coring (APC), then half-length APC, and then extended core barrel (XCB) coring as necessary. Temperature measurements used the advanced piston corer temperature (APCT-3) and Sediment Temperature 2 (SET2) tools (Neumann et al. 2023). Downhole logging conducted after coring used the triple combination and Formation MicroScanner sonic logging tool strings. After bringing core sections onto the core receiving platform of the D/V *JOIDES Resolution*, whole round samples for microbiology were retrieved within ∼30 minutes using ethanol-cleaned spatulas. Whole round samples for DNA-based studies were capped with ethanol-sterilized endcaps, transferred to the microbiology laboratory, and stored briefly at 4°C in heat-sealed tri-foil gas-tight laminated bags flushed with nitrogen until processing. Masks, gloves and laboratory coats were worn during sample handling in the laboratory where core samples were transferred from their gas-tight bags onto sterilized foil on the bench surface inside a Table KOACH T 500-F system, which creates an ISO Class I clean air environment (Koken Ltd., Japan). In addition, the bench surface was targeted with a fanless ionizer (Winstat BF2MA, Shishido Electrostatic Co., Ltd., Japan). Within this clean space, the exterior 2 cm of the extruded core section were removed using a sterilized ceramic knife. The core interior was transferred to sterile 50-mL Falcon tubes, labeled, and immediately frozen at −80°C for post cruise analyses. For RNA-based studies, sampling occurred immediately after core retrieval on the core receiving platform by sub-coring with a sterile, cutoff 50cc syringe into the center of each freshly cut core section targeted. These sub-cores were immediately frozen in liquid nitrogen and stored at −80°C.

### DNA extraction and sequencing

DNA was extracted from selected core samples using a FastDNA SPIN Kit for Soil (MP Biomedicals). Up to 5 grams of sediment were processed following the manufacturer’s protocol with homogenization modifications as described previously (Ramírez et al., 2018). Briefly, sediment was homogenized twice in Lysing Matrix E tubes for 40s at speed 5.5, with a 2-minute pause on ice between cycles. Extracts were concentrated using Amicon Ultra-0.5 ml Centrifugal Filters (MilliporeSigma). A control extraction, in which no sediment was added, was included to account for any laboratory contaminants. Thirteen libraries for metagenome sequencing were prepared from genomic DNA extracts at the University of Delaware DNA Sequencing & Genotyping Center and sequenced with Illumina NovaSeq S4 PE150 chemistry at the University of California, Davis Genome Center. Metagenome sequence reads were deposited to the National Center for Biotechnology Information Sequence Read Archive under access numbers SRR580794-SRR580807.

### Metagenomic co-assembly, binning, and taxonomic assignment

Metagenomes originating from adjacent regions (such as adjacent depths targeted in this study) are likely to overlap in the sequence space, increasing the mean coverage and extent of reconstruction of MAGs when using a co-assembly approach. For MAG reconstructions, the trimmed reads of metagenomic datasets from all 13 Guaymas samples sequenced in this study, as well as 13 additional Guaymas metagenomes from IODP385 that were previously sequenced at the University of Delaware DNA Sequencing & Genotyping Center (Bojanova et al., in review), were co-assembled into contigs using MEGAHIT 1.2.9 (Li et al., 2016) with default parameters. Assembled contigs were binned using MetaBAT2 2.12.183 (Kang et al., 2015), MaxBin2 2.2.7 (Wu et al., 2016), as well as CONCOCT 1.1.0 (Alneberg et al., 2013), and output bins from all three binning algorithms were refined and dereplicated using DAS Tool 1.1.6 (Sieber et al., 2018). Completeness, size, and contamination levels of the reconstructed genomes were estimated using CheckM2 1.0.0 (Chklovski et al., 2022). Only MAGs that were at least 50% complete and contained less than 10% contamination were used for downstream analyses (Supplementary Table 1). The taxonomic placement of the MAGs was performed with GTDB-Tk 2.1.1 (Chaumeil et al., 2020). Sequencing control samples identified MAGs of lab/control contaminants, including Patescibacteria (classes Paceibacteria and Microgenomatia), Actinobacteriota (classes Actinomycetia and Humimicrobia), Gammaproteobacteria (orders Pseudomonadales and Burkholderiales), and Firmicutes (order Staphylococcales); these were removed from downstream analyses (Supplementary Table 2).

### Calculation of MAG relative abundances

Metagenomic reads from 26 samples were mapped to each MAG using the coverM 0.6.1 (https://github.com/wwood/CoverM) command line tool with the BWA 2.0 aligner (Vasimuddin et al., 2019). The CoverM tool automatically concatenated all the MAGs into a single file, and metagenomic reads were recruited to MAG contigs, setting the parameter --min-read-percent-identity to 95 and --min-read-aligned-percent to 50. The “Relative Abundance” CoverM method on the “genome” setting was used to calculate the percent of total metagenomic reads per sample that mapped to each of the 89 MAGs. A custom R script was utilized to concatenate all coverM output files into a single file in a matrix format (with each sample representing a column and each row representing total percent of DNA-Seq reads per sample that mapped to a MAG) and was used for heatmap plotting.

### Gene annotation, and prediction of KEGG metabolic module presence/absence using MetaPathPredict

Genes were called for all MAGs using Prodigal 2.6.3 (Hyatt et al., 2010) and then annotated using Prokka 1.14.6 (Seemann, 2014) KofamScan 1.3.0 (Aramaki et al., 2020), and METABOLIC 4.0 (Zhou et al., 2022) using default settings. KEGG modules for bacterial MAGs were reconstructed using gene annotations from the KofamScan 1.3.0 (Aramaki et al., 2020) command line tool, and the presence or absence of incomplete modules in the genomes was predicted using MetaPathPredict 1.0.0 (Geller-McGrath et al., 2022) with default settings. MetaPathPredict cannot yet be applied to archaeal MAGs. Briefly, Prodigal was used to call genes, and KofamScan (Aramaki et al., 2020) was used to annotate them. Gene annotations were generated for predicted genes from bacterial MAGs, and were used as input to MetaPathPredict, which generated predictions for the presence or absence of KEGG modules based on the gene annotations of all bacterial MAGs.

### nMDS analysis of MAG abundances in metagenomic datasets and associated environmental parameters

The abundances of metagenomic reads mapped to MAG contigs were normalized as transcripts per million using coverM (https://github.com/wwood/CoverM), analyzed using nMDS, and fitted with all environmental data in Supplementary Table 1 in R (Team, 2018) using the metaMDS and envfit functions from the vegan 2.6-4 package (Dixon, 2003). The results were plotted using ggplot2 3.3.6 (Wickham et al., 2016) with sample shapes corresponding to temperature regime in Figure 4, and depth regime in Supplementary Figure 4.

### Estimated genome size analysis

The estimated genome size of all 89 MAGs was calculated by dividing the MAG assembly size (total base pair length of the MAG) by the fractional CheckM2 completeness of the MAG (the default CheckM2 completeness output divided by 100; a number between 0 and 1). Difference in genome size distributions for MAGs that recruited at least 0.1% of metagenomic reads from samples across temperature (cool [2-20°C], warm [20-45°C], hot [45-60°C]) and depth (shallow [2-15 mbsf], intermediate [15-60 mbsf], deep [>60 mbsf]) regimes was assessed using the two-sided Welch’s *t*-test, and resulting *p*-values were adjusted for multiple comparisons via Benjamini-Hochberg correction. The average estimated genome size of MAGs that recruited at least 0.1% of reads from metagenomic samples (*n* = 26) was fitted using linear regression against temperature and depth measurements affiliated with the samples.

### MAG recovery at sampled sites versus temperature and depth

The number of unique MAGs that recruited at least 0.1% of reads from metagenomic samples (*n* = 26) was fitted using linear regression against temperature and depth measurements affiliated with the samples.

### Scanning of MAGs for secondary metabolite biosynthetic gene clusters

All 89 MAGs were individually scanned for secondary metabolic biosynthetic gene clusters using antiSMASH 6.0 (Blin et al., 2021) with default parameters. Resulting gene cluster prediction results (in GenBank format) were parsed and their gene content was analyzed. Clusters with a total length less than 5kb were discarded from downstream analysis to minimize the inclusion of fragmented biosynthetic clusters in the analysis.

### RNA extraction, sequencing, and mapping of RNA reads to the MAGs

RNA was extracted from 19 sediment samples from sites U1545B-U1552B and a blank sample (control) using the RNeasy PowerSoil Total RNA Kit (Qiagen) following the manufacturer’s protocol. RNA samples were prepared from samples spanning the depths 0.8 to 101.9 mbsf. All samples, including a blank control, were first washed twice with absolute ethanol (200 proof; purity ≥ 99.5%), and sterile DEPC water (once) to reduce hydrocarbons and other inhibitory elements that otherwise resulted in low RNA yield. In brief, 13-15 grams of frozen sediments were transferred into UV-sterilized 50 ml Falcon tubes (RNAase/DNase free) using clean, autoclaved and ethanol-washed metallic spatulas. Each sample transferred into the 50 ml Falcon tube received an equal volume of absolute ethanol and was shaken manually for 2 min followed by 30 seconds of vortexing at full speed to create a slurry. Samples were spun in an Eppendorf centrifuge (5810R) for 2 minutes at 2000 rpm. The supernatant was decanted and after the second wash with absolute ethanol, an equal volume of DEPC water was added into each sample and samples were spun for 2 minutes at 2000 rpm. The supernatant was decanted, and each sediment sample was immediately divided into three 15 mL Falcon tubes containing beads provided in the PowerSoil Total RNA Isolation Kit (Qiagen). The RNA extraction protocol was followed as suggested by the manufacturer, with the modification that the RNA extracted from the three aliquots was pooled into one RNA collection column. All steps were performed in a UV-sterilized clean hood equipped with HEPA filters. Surfaces inside the hood and pipettes were thoroughly cleaned with RNase AWAY^TM^ (Thermo Scientific™) before every RNA extraction and in between extraction steps.

Trace DNA contaminants were removed from RNA extracts using TURBO DNase (Thermo Fisher Scientific) and the manufacturer’s protocol. Removal of DNA was confirmed by negative PCR reactions using the bacterial primers BACT1369F/PROK1541R (F: 5’CGGTGAATACGTTCYCGG 3’, R: 5’AAGGAGGTGATCCRGCCGCA 3’; Suzuki et al., 2000), targeting the 16S rRNA gene. Each 25 μl PCR reaction was prepared using 0.5 U μl^−1^ GoTaq® G2 Flexi DNA Polymerase (Promega), 1X Colorless GoTaq® Flexi Buffer, 2.5 mM MgCl_2_, (Promega) 0.4 mM dNTP Mix (Promega), 4 μM of each primer (final concentrations), and DEPC water. PCR reactions used an Eppendorf Mastercycler Pro S Vapoprotect (Model 6321) thermocycler with following conditions: 94°C for 5 min, followed by 35 cycles of 94°C (30 s), 55°C (30 s), and 72°C (45 s). The PCR products were run in 2% agarose gels (Low-EEO/Multi-Purpose/Molecular Biology Grade Fisher BioReagents™) to confirm absence of DNA amplification. Amplified cDNAs were prepared from DNA-free RNA extracts using the Ovation RNA-Seq System V2 (NuGEN, San Carlos, CA). Libraries were sequenced on Illumina NextSeq 500 PE 150 High Output at the Georgia Genomics and Bioinformatics Core. The cDNA library generated from our control did not contain detectable DNA. It was nonetheless submitted for sequencing, but it failed to generate any sequences that met the minimum length criterion of 300-400 base pairs.

Reads from the 13 metatranscriptome samples collected from sites that metagenomic samples were taken from were mapped to each MAG using the CoverM 0.6.1 (https://github.com/wwood/CoverM) command line tool with the BWA 2.0 aligner (Vasimuddin et al., 2019). The CoverM tool automatically concatenated all the MAGs into a single file, and metatranscriptome reads were recruited to MAG contigs, setting the parameter --min-read-percent-identity to 95 and --min-read-aligned-percent to 50. A custom R script was utilized to concatenate all coverM output files into a single file in a matrix format, with each sample representing a column and each row representing total percent of RNA-Seq reads per sample that mapped to a MAG. The output was used in this study for heatmap plotting to examine evidence for activity of the taxa for which we recovered MAGs. Metatranscriptome reads were deposited to the National Center for Biotechnology Information Sequence Read Archive under accession numbers SRR22580929-SRR22580947.

### Cell counts

The sediment sampling for cell counts occurred immediately after core retrieval on the core receiving platform by sub-coring with a sterile, tip-cut 2.5 cc syringe from the center of each freshly cut core section. Approximately 2 cm^3^ sub-cores were immediately put into tubes containing fixation solution consisting of 8 mL of 3xPBS (Gibco™ PBS, pH 7.4, Fischer) and 5% (v/v) neutralized formalin (Thermo Scientific™ Shandon™ Formal-Fixx™ Neutral Buffered Formalin). If necessary, the mixture was stored at 4°C.

Fixed cells were separated from the slurry using ultrasonication and density gradient centrifugation (Morono et al., 2013). For cell detachment, a 1 mL aliquot of the formalin-fixed sediment slurry was amended with 1.4 mL of 2.5% NaCl, 300 μL of pure methanol, and 300 μL of detergent mix (Kallmeyer et al. 2008), 100 mM ethylenediamine tetraacetic acid [EDTA], 100 mM sodium pyrophosphate, 1% [v/v] Tween-80). The mixture was thoroughly shaken for 60 min (Shake Master, Bio Medical Science, Japan), and subsequently sonicated at 160 W for 30 s for 10 cycles (Bioruptor UCD-250HSA; Cosmo Bio, Japan). The detached cells were recovered by centrifugation based on the density difference of microbial cells and sediment particles, which allows collection of microbial cells in a low-density layer. The sample was transferred onto a set of four density layers composed of 30% Nycodenz (1.15 g cm^−3^), 50% Nycodenz (1.25 g cm^−3^), 80% Nycodenz (1.42 g cm^−3^), and 67% sodium polytungstate (2.08 g cm^−3^). Cells and sediment particles were separated by centrifugation at 10,000 × g for 1 h at 25°C. The light density layer was collected using a 20G needle syringe. The heavy fraction, including precipitated sediment particles, was resuspended with 5 mL of 2.5% NaCl, and centrifuged at 5000 × g for 15 min at 25°C. The supernatant was combined with the previously recovered light density fraction. With the remaining sediment pellet, the density separation was repeated. The sediment was resuspended using 2.1 mL of 2.5% NaCl, 300 μL of methanol, and 300 μL of detergent mix and shaken at 500 rpm for 60 min at 25°C, before the slurry sample was transferred into a fresh centrifugation tube where it was layered onto another density gradient and separated by centrifugation just as before. The light density layer was collected using a 20G needle syringe, and combined with the previously collected light density fraction and supernatant to form a single suspension for cell counting.

For cell enumeration, a 50%-aliquot of the collected cell suspension was passed through a 0.22-μm polycarbonate membrane filter. Cells on the membrane filter were treated with SYBR Green I nucleic acid staining solution (1/40 of the stock concentration of SYBR Green I diluted in Tris-EDTA [TE] buffer). The number of SYBR Green I– stained cells were enumerated either by a direct microscopic count (Inagaki et al., 2015) or an image-based discriminative count (Morono et al., 2009). For image-based discriminative counting, the Count Nuclei function of the MetaMorph software (Molecular Devices) was used to detect and enumerate microbial cells.

## Data availability

All metagenome and metatranscriptome data discussed in this manuscript are deposited to NCBI GenBank under SRR22580929-SRR22580947. All custom scripts used for data analysis and figure creation are available in the following GitHub repository: https://github.com/d-mcgrath/guaymas_basin. All environmental data are available through BCO-DMO (https://www.bco-dmo.org/project/833856) and the IODP Expedition 385 report (http://publications.iodp.org/proceedings/385/385title.html)

## Acknowledgments

The authors would like to acknowledge the crew and entire science party for IODP Expedition 385 for their assistance with sample collection. Without their assistance this study would be impossible. The authors would also like to thank Gustavo Ramírez for his assistance with DNA extractions using his method. We thank Mark Shaw and Bruce Kingham in the University of Delaware DNA Sequencing & Genotyping Center for assistance with sample preparation and Illumina sequencing. This study was supported by NSF Grant OCE-2046799 to VE, PM, AT, and R. Hatzenpichler, by NSF grant OCE-1829903 to VE, PM, and AT, and by JSPS KAKENHI Grants JP19H00730 and JP23H00154 to YM.

## Author contributions

VE, AT, and PM designed the study. VE took primary responsibility for collecting the samples during IODP Expedition 385. AT served as Co-Chief Scientist of IODP Expedition 385. DB and PM extracted DNA and RNA for metagenomes and metatranscriptomes, respectively, and DB prepared metagenome libraries for sequencing, and handled data deposition into GenBank. DGM took primary responsibility for bioinformatic processing of metagenome data and mapping of transcripts to MAGs. DGM and PM analyzed the MAG data and VE and AT contributed to data interpretation. YM provided cell count data. DGM, PM, VE, and AT co-wrote the first draft of the manuscript, and all authors contributed to its final form.

## Competing interests

The authors declare no competing interests.

## Notes

https://github.com/d-mcgrath/guaymas_basin

### Competing Interest Statement

The authors have declared no competing interest.

