## Supplementary Text and Supplementary Figures for "Metagenomic Profiles of Archaea and Bacteria within Thermal and Geochemical Gradients of the Guaymas Basin Deep Subsurface"

**Supplementary Material**

**Extended overview on bacterial and archaeal lineages in Guaymas Basin**

**Chloroflexota.** Genes detected in our Chloroflexota MAGs are associated with fatty acid degradation, ornithine biosynthesis and the methionine salvage pathway that produces methionine by recycling sulfur-bearing metabolites (Sekowska et al., 2004). Ornithine biosynthesis was also evident in MAGs affiliated with other bacterial and archaeal phyla, including the Hadarchaeota. Genes for enzymes involved in ornithine synthesis (e.g., ornithine decarboxylase) have been documented previously in deep biosphere samples (Orsi et al., 2013), indicating that this capability may be widely utilized by subsurface microbiota (Hernández et al., 2021).

The majority of Chloroflexota genomes contained marker genes associated with the oxidation of formate (*fdhA*, *fdhB*, *fdoG*, *fdoH*; McGonigle et al., 2020) and carbon monoxide (*coxL*, *coxM*, *coxS*; Islam et al., 2019). Their occurrence coincides with increased carbon monoxide concentrations (83-665 nM) between 0.8-60 mbsf (Supplementary Table 1). Hydrogen utilization is suggested by Ni-Fe hydrogenases and key genes of acetate/acetyl-CoA production (*pta*/*ack*, *acdA*) in Chloroflexota MAGs (Supplementary Tables 4-5). Hydrogen concentrations associated with our metagenomic samples ranged between 23-84 nM (Supplementary Table 1) suggesting active hydrogen cycling (Lin et al., 2012). Genes associated with the Wood-Ljungdahl pathway (WL; *cdhD*, *cdhE*, *cooS*) were present in 17/23 Chloroflexota MAGs, while ATP-citrate lyase (*aclA*), associated with the reductive TCA cycle (rTCA), was present in one MAG from the VGOG01 order. *mmoB* gene associated with methane oxidation was identified in two Chloroflexota MAGs from the orders Promineofilales (Speirs et al., 2019) and E44-bin15 (associated with petroleum seepage in marine sediments; Supplementary Table 6; Dong et al., 2019). Previous DNA-SIP experiments identified members of the Chloroflexota as putative methane oxidizers and detected methane monooxygenases (*mmoX* or *pmoB)* in Chloroflexota genomes (Altshuler et al., 2022). We caution that the identified *mmoB* has a regulatory role in methane oxidation, while the catalytic activity is encoded by the *mmoH* gene in the *mmo* operon (Sirajuddin & Rosenzweig, 2015).

Genes involved in sulfate and sulfur assimilation for amino acid synthesis (e.g., cysteine; *cysN*, *sat*, *cysD*) and in sulfur oxidation *(sdo*, *dsrH)* were present in all Chloroflexota MAGs, however, none of our Chloroflexota MAGs encoded complete pathways for sulfur oxidation. Evidence for dimethyl sulfoxide (DMSO) utilization was suggested by the presence of *dmsA* and/or *dmsB* (involved in DMSO reduction) in five of our MAGs (Supplementary Tables 4-5). DMSO is abundant in deep sea ecosystems and was suggested to be an electron acceptor for microbes that survive in deep-sea extreme conditions (Xiong et al., 2016). Genes for denitrification, (e.g., *nirB* and *nirD*) were encoded in one Chloroflexota MAG. Finally, genes for transport of tungstate were identified in 4 Chloroflexota MAGs, and transporters of molybdenum/tungsten in 10 MAGs (Supplementary Tables 4-5). Tungsten is found abundantly in hydrothermal ecosystems (Kishida et al., 2004), and serves as a redox catalyst in metalloenzymes of thermophilic archaea inhabiting hydrothermal vents (Kletzin & Adams, 1996) and hot springs (Buessecker et al., 2022). Evidence of putative arsenate biomineralization for detoxification (e.g., *arsC*, *arsM* genes) or energy gain via arsenotrophy (*arrA* gene; Saunders et al., 2019) was present in three MAGs. Sedimentary arsenic input would explain the elevated Arsenic concentrations in Guaymas Basin hydrothermal fluids (up to 1 mmol) that exceed those at other hydrothermal vent sites (Von Damm et al., 1985). Five Chloroflexota MAGs contained CRISPR/Cas genes involved in genome editing (Supplementary Table 4-5).

**Thermoproteota.** Bathyarchaeia recycle hydrogen and CO_2_ from fermentation using the WL pathway (He et al., 2016). Eight out of 11 of our Bathyarchaeia MAGs contained genes involved of the WL pathway. In addition, we detected the marker gene for the formaldehyde activating enzyme (*fae*) in 5 MAGs affiliated with 40CM-2-53-6, B26-1 and TCS64 orders. *Fae* condenses formaldehyde and tetrahydromethanopterin to form methylenetetrahydromethanopterin that can be reduced and utilized in the WL pathway (Timmers et al., 2017; Vorholt et al., 2000). Six Bathyarchaeia MAGs encoded the *mer* gene for the reduction of methylenetetrahydromethanopterin in the WL pathway. Various Bathyarchaeia sub-lineages have been reported to encode genes for anaerobic oxidation of methane/alkane compounds (*mcr*/*acr* complex) (Evans et al., 2015, 2019; Qi et al., 2021; Vanwonterghem et al., 2016). We detected the methane/alkane oxidation marker genes *fwd*, *ftr*, *mtd*, *mch*, and *mtr* in all of our Bathyarchaeia MAGs. However, the *acr*/*mcr* genes encoding the methyl/alkyl-coenzyme M complex were absent, indicating loss of the *mcr*/*acr* operon as described previously for Bathyarchaeia . Nine out of 11 Bathyarchaeia MAGs also encoded genes for acetate formation (*acdA*, *ack*, *pta, acs)* which could be utilized to couple methylotrophy with acetogenesis, as has been described previously in Bathyarchaeia MAGs from deep sediments (Farag et al., 2020; He et al., 2016).

Marker genes for other specific metabolic capacities in our Bathyarchaeota MAGs included genes for fermentation (*porA*), hydrogen cycling (Ni-Fe hydrogenases) and genes involved in the anaerobic degradation of benzoate (*bcrA, bcrB, bcrD*; Kung et al., 2009). The *bcr* genes were also observed in Chloroflexota (5 MAGs), Zixibacteria (1 MAG) and Desulfobacterota (4 MAGs); the latter group is easily enriched from Guaymas Basin sediments on benzoate under sulfate-reducing conditions (Edgcomb et al., 2022).

**Acidobacteriota.** Members of the highly diverse heterotrophic phylum Acidobacteriota occur in a wide range of freshwater and marine seafloor environments and can utilize oxygen or other electron acceptors for respiration (e.g., nitrate, nitrite, sulfate) (Flieder et al., 2021 and references therein). We detected eight Acidobacteriota MAGs annotated to the orders of Aminicenantales (7 MAGs), and Acidoferrales (1 MAG). Aminicenantales MAGs were previously recovered from surficial Guaymas Basin sediments (Dombrowski et al., 2018), and in this study they were found at all sites between 0.8-60 mbsf. The Acidoferrales MAG was detected only below 60 mbsf at sites U1545 and U1548. The overall metabolic potential of Guaymas subsurface Acidobacteriota MAGs is described in Supplementary Tables 4 and 5.

We observed that almost all Acidobacteriota MAGs encoded the NtrY-NtrX two-component regulatory system which can be important in the Guaymas subsurface. NtrY-NtrX is a redox sensor system widely distributed in Proteobacteria which regulates denitrification and nitrogen fixation genes (Pawlowski et al., 1991; DelVecchio et al., 2002), and senses nitrogen levels under nitrogen limitation (Bonato et al., 2016). Six of our eight Acidobacteriota MAGs contained at least one gene of the *nif* operon (e.g., *nifU, nifB, nifH*) which suggests putative nitrogen fixation in our subsurface samples, similar to previous reports of deep-sea sediment Acidobacteriota that encode *nifH* in their genomes (Kapili et al., 2020). Two Acidobacteriota MAGs contained CRISPR/Cas genes involved in genome editing (Supplementary Table 4).

**Desulfobacterota.** Desulfobacterota include primarily sulfate reduces and syntrophic lineages that couple sulfate reduction with methane and short-chain alkane oxidation by anaerobic methane oxidizers (ANME archaea). These bacteria are widespread in Guaymas Basin sediments and other hydrothermal and cold seep sites (Knittel & Boetius, 2009; Murphy et al., 2021; Speth et al., 2022; Wegener et al., 2022; Zhou et al., 2022). Seven Desulfobacterota MAGs, belonging to the Desulfobacterales, Desulfatiglandales, and WTBG01 and WVXP0 orders were recovered primarily from shallow sulfate-rich cool sediments of all sites (0.8-15 mbsf, at or above the SMTZ with temperatures 2-20^o^C) (Supplementary Figure 4). Two Desulfobacterales MAGs contained the *dsr* operon (e.g., *dsrB/J/K/D*) involved in dissimilatory sulfate reduction (Venceslau et al., 2014). One MAG annotated to the WTBG01 order (found in freshwater anoxic sulfidic sediments; Murphy et al., 2021) encoded the sulfate adenylyltransferase gene (*sat*) associated primarily with sulfur assimilation. Marker genes of DMSO reduction and/or sulfur assimilation were also detected in all Desulfobacterota MAGs. One Desulfobacterota MAG contained CRISPR/Cas genes involved in genome editing (Supplementary Table 4).

The potential for iron reduction was evidenced in all Desulfobacterota MAGs by the presence of *mtrA*, *mtoA* (Garber et al., 2020) and *eetB* genes, suggesting an extracellular electron transfer mechanism. In addition, *DFE* genes encoding multiheme cytochromes (e.g., *DFE_0449, DFE_0461, DFE_0451*) were found in three of our MAGs annotated to Desulfobacterota and in five MAGs annotated to Aminicenantales (Acidobacteriota). *DFE* genes are involved in iron oxidation and were originally documented in the genome of *Desulfovibrio ferrophilus* strain IS5 (Deng & Okamoto, 2018). They encode cytochromes and β-propeller proteins, which can function as electron carriers and leader peptides in extracellular electron transfer (Chatterjee et al., 2021; Deng & Okamoto, 2018). At depths where Desulfobacterota MAGs were detected, dissolved porewater iron ranged in concentration from less than 1 μM to greater than 4 μM (Supplementary Table 1), indicating possibly active iron cycling with little accumulation.

**Aerophobota and White Oak River group 3 (WOR-3).** The phylum Aerophobota is widely distributed in deep-sea sediments, and includes fermentative thermophiles and hyperthermophiles affiliated with hydrocarbon-rich environments, and hydrothermal sites including Guaymas Basin (Speth et al., 2022). Members of the Aerophobota are also abundantly found in hydrate-bearing sediments (Liu et al., 2022). We recovered five Aerophobota MAGs (order Aerophobiales) from the deep subsurface at depths below 100 mbsf, as long as temperatures did not exceed 40°C. Thus, the distribution of Aerophobota appears to be constrained by thermal limits rather than by depth. Guaymas Aerophobota MAGs encoded marker genes for acetate/acetyl-CoA production (*acdA*, *ack*, *pta*) as previously described (Dong et al., 2019), fermentation (*porA*) and degradation of polysaccharides (e.g., cellulose, chitin). Two Aerophobota MAGs contained CRISPR/Cas and CRISPR/Csm genes that comprise adaptive defense systems against infectious agents in prokaryotes (Colognori et al., 2023).

Members of the bacterial WOR-3 candidate phylum were originally described from estuarine (Baker et al., 2015) and hydrothermal Guaymas Basin (Dombrowski et al., 2017) sediments. Five of our six WOR-3 MAGs were from depths at 0.8-26.9 mbsf, while the only WOR-3 MAG from the UBA3073 order was recovered primarily from 112.5 and 154.2 mbsf at site U1545 (up to ~45°C) (Figure 2). Our WOR-3 MAGs encoded various peptidases, as well as genes for H_2_ cycling (Ni-Fe hydrogenase genes) and putative chitin degradation (endo-acting chitinase genes). While these results match previous findings (Baker et al., 2015), the Guaymas subsurface WOR-3 MAGs also contain genes for fermentation and acetate production (*acs, acdA*, *ack, porA*), and marker genes for endohemicellulases and amylolytic enzymes that can degrade other polysaccharides aside from chitin. One WOR-3 MAG encoded CRISPR/Cas genes.

**Iron reduction and oxidation**. Iron reduction is a known capability for Bacteria and Archaea associated with marine seafloor sediments (Flieder et al., 2021; Jiang et al., 2019). Marker genes involved in dissimilatory iron reduction (e.g., *dmkA*, *dmkB*, *eetA*, *eetB*, *fmnA*, *fmnB*, *pplA*, *ndh2*; Garber et al., 2020) with the potential for extracellular electron transfer (EET) were identified in 86/89 MAGs from all recovered phyla (Supplementary Tables 4-5), and were only missing from one Thermoproteota, one WOR-3, and one Aenigmatarchaeota MAG. These genes participate in EET from the cell towards the surrounding environment (Light et al., 2018; Shi et al., 2016). Based on laboratory experiments, EET is suggested to enhance iron bioavailability and iron uptake in anaerobes, and to act as a redox mechanism that may contribute to the proton motive force (Jeuken et al., 2020).

Eight out of 23 Chloroflexota, 2/11 Thermoproteota, 5/8 Acidobacteriota, 3/7 Desulfobacterota, and 1/6 WOR-3 MAGs also encoded components of the *DFE_0448*-*0451* and *DFE_0461*-*0465* operon genes (Supplementary Tables 4-5) homologous to multiheme cytochrome systems first identified in *Desulfovibrio ferrophilus* (Deng et al., 2018). According to the *D*. *ferrophilus* model, electrons from an external iron source move along extracellular and membrane-spanning multiheme cytochromes from the outer membrane to the periplasm of the cell and finally are passed to a terminal electron acceptor (Deng et al., 2018). Putative terminal electron acceptors for this process in MAGs containing *DFE* components included the sulfur cycle intermediates sulfite (based on the presence of sulfite reductase *asrA* and *asrB*), tetrathionate (from the detection of tetrathionate reductase gene *ttrB*), and thiosulfate (due to the annotation of thiosulfate reductase *phsA/B* genes; Supplementary Tables 4-5). The capacity for polysulfide reduction was also detected based on the presence of polysulfide reductase (*psrA*) in four Acidobacteriota MAGs containing *DFE* multiheme cytochrome components.

Electron acceptors from nitrogen cycle intermediates included nitrite (*nasD*) and nitrate (*napA*, *narB*). This process is predicted to be utilized when more energy-rich substrates such as organic compounds are limited (Deng et al., 2018). Iron (II and III) concentrations associated with metagenomic samples containing MAGs that encoded genes affiliated with iron metabolism ranged from 0 to 11.8 µM (Supplementary Table 1).

**Carbon monoxide oxidation**. Hydrothermal environments often contain carbon monoxide (CO), which can be produced by the breakdown of organic matter or generated by certain anaerobic microorganisms (Kochetkova et al., 2011; Sokolova et al., 2009). The potential for CO oxidation, an energetically favorable anaerobic reaction, is prevalent in subsurface bacterial and archaeal MAGs (Baker et al., 2016; Magnabosco et al., 2016). Nine out of 23 Chloroflexota, 3/11 Thermoproteota, 7/8 Acidobacteriota, 3/7 Desulfobacterota, 2/5 Aerophobota, and one Hadarchaeota MAG contained the genes *coxM* and *coxS* encoding CO dehydrogenase subunits (Supplementary Tables 4-5). Genes for catalytic nickel-containing CO dehydrogenase (*cooS*) and/or its iron-sulfur subunits (*cooF*) were further identified in 9/23 Chloroflexota, 1/11 Thermoproteota, 1/8 Acidobacteriota, 3/7 Desulfobacterota, and 2/5 Aerophobota. The presence of these genes suggests the capacity to oxidize CO and H_2_O to CO_2_ and H_2_. Bacteria and Archaea are also capable of CO oxidation coupled to the anaerobic reduction of sulfur and nitrogen compounds (King, 2006; Oelgeschläger & Rother, 2008). Potential terminal electron acceptors for our CO-oxidizing MAGs included the sulfur cycle intermediates sulfite (due to the presence of genes *asrA* and *asrB*), tetrathionate (*ttrB*), thiosulfate (*phsA/B*), and polysulfide (*psrA*; Supplementary Tables 4-5). CO concentrations associated with metagenomic samples containing MAGs that encoded these genes ranged from 83 to 665 nM (Supplementary Table 1).

**MetaPathPredict insights into metabolic potential of Zixibacteria and Cloacimonadetes.** Some phyla for which we recovered MAGs (e.g., Zixibacteria and Cloacimonadota) remain poorly described in terms of their metabolic potential. To assess and predict the metabolic potential of some of our less-complete and poorly characterized bacterial MAGs, we applied a new tool “MetaPathPredict” (Geller-McGrath et al., 2022) to our Zixibacteria and Cloacimonadota MAGs. MetaPathPredict is a software designed to predict the presence or absence of complete KEGG modules in partially complete bacterial genomes (see below). The tool utilizes machine learning models trained on bacterial gene annotations in the format of KEGG gene orthologs to predict the presence or absence of whole KEGG modules, and has been designed to handle gene annotations from incomplete bacterial genomes. It is very common in environmental metagenomic studies that reconstructed MAGs will vary in their degree of completeness. Usually only a fraction exceeds > 80% completeness. Application of MetaPathPredict to such partially complete MAGs can be useful for providing predictions on whether metabolic pathways are present especially in cases where some key genes are missing. While this tool can sometimes miss predictions of pathways that should be present based on benchmarking tests with genomic data (Geller-McGrath et al., 2022), it can nonetheless be useful for gaining insights into less-complete MAGs, and also for predicting metabolic capacities of MAGs affiliated with poorly understood taxonomic groups.

Our Zixibacteria MAGs were 61-88.5% complete and our single Cloacimonadota MAG was 75% complete. The results of the MetaPathPredict analysis are described below and presented in Supplementary Table 7 which shows: 1) KEGG modules present and complete in our Zixibacteria and Cloacimonadota MAGs, and predicted by MetaPathPredict 2) KEGG modules absent or incomplete but predicted to be present by MetaPathPredict, and 3) incomplete KEGG modules, also predicted to be absent by MetaPathPredict.

Lineages of Zixibacteria have been documented in various marine and terrestrial subsurface ecosystems, hypersaline settings and anoxic sediments (Anantharaman et al., 2018; Baker et al., 2015; Castelle et al., 2013; Lin et al., 2012b; Momper et al., 2017; Wong et al., 2020). This taxon is thought to be capable of dissimilatory nitrate and sulfate reduction, and it seems to lack complete carbon fixation pathways (Momper et al., 2017). The 4 Zixibacteria MAGs we recovered were detected at all sampling sites down to 25.8 mbsf and metagenome reads mapped most intensely from shallow (8.6-16.2 mbsf) depths. Zixibacteria (order MSB-5A5) have the metabolic capacity for oxidation of fatty acids, (e.g., acyl-CoA dehydrogenase) and synthesis of a suite of vitamins (e.g, B6, B1). MetaPathPredict predicted the potential for dissimilatory sulfate reduction in one of our MAGs (order DG-27), which also contained a marker gene for this process (*dsrA*). Two Zixibacteria MAGs (orders DG-27 and UBA10806) were also predicted to encode the potential for dissimilatory nitrate reduction and contained marker genes *napB* and *nrfH*. MetaPathPredict additionally predicted the synthesis of vitamin B7 in two Zixibacteria MAGs, and acetate production via the phosphate acetyltransferase-acetate kinase (Pta-Ack) pathway in all four MAGs. The Pta-Ack pathway can produce acetate/acetyl-CoA and can participate in carbon fixation by providing acetyl-CoA. Complete acetogenesis via the pta-ack pathway was also confirmed from our genomic data in two MAGs (*ack*, *pta*). MetaPathPredict predicted various transporters (e.g., phosphate, iron and ABC transporters), pathways involved in synthesis of co-factors using amino acids or tRNA reductases (e.g., from glutamate to heme; from tRNA-glutamyl to sideroheme), and the synthesis of various amino acids (e.g., threonine, serine, valine isoleucine). Many of these processes (e.g., biosynthesis of amino acids, iron reduction) were also verified with marker genes. None of the Zixibacteria MAGs were predicted to contain the potential for carbon fixation.

Cloacimonadota are abundant in anoxic/sulfidic water columns and cold seep brine pools (e.g., Black and Red Sea, respectively; Suominen et al., 2021; Villanueva et al., 2021; Zhang et al., 2016). They are suggested to perform diverse metabolisms including, carbon fixation, fermentation, and assimilation of proteins as carbon and nitrogen sources. Our single Cloacimonadota MAG was present in low abundance at site U1545B in metagenomes between 25.8 and 63.8 mbsf. The Cloacimonadota MAG was not predicted to contain the capacity for carbon fixation, however our data showed (and the MetaPathPredict successfully predicted) that this taxon encodes genes for fermentation via acetate production (*pta*, *ack*), and synthesis of vitamin B7 and salvage of thiamine (vitamin B1), an indispensable cofactor in amino acid and carbohydrate metabolism; salvage of B1 is linked to the metabolism of pyrimidines (Gonçalves & Gonçalves, 2019). Biosynthesis of purines and pyrimidines is a core metabolic process, and was detected, and predicted, in our Cloacimonadota MAG.

**Supplementary Figures**

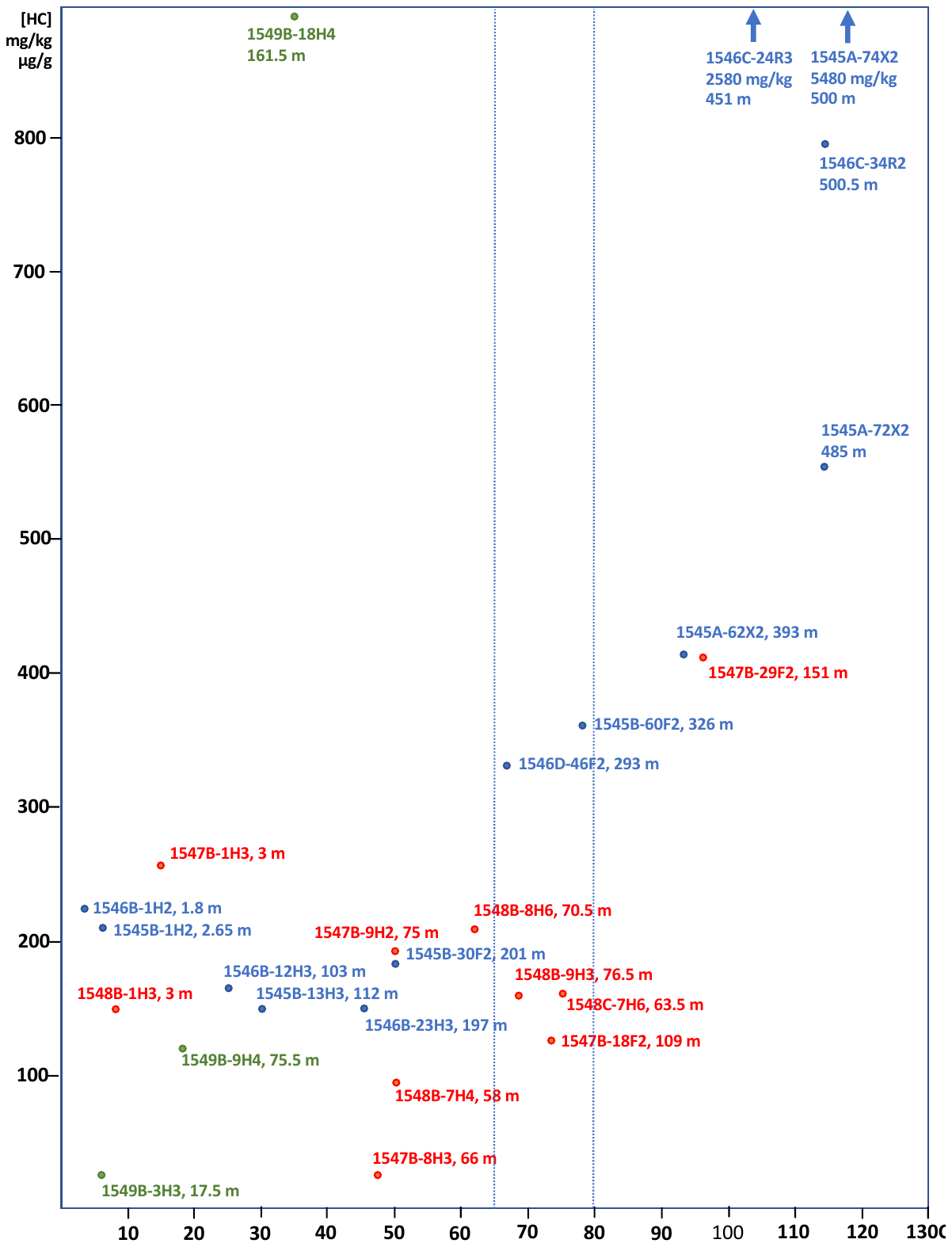

**Supplementary Figure 1**. **Total Hydrocarbon content of Guaymas Basin sediments**. Samples were analyzed at Alpha Analytical (Mansfield, MA, USA) for fingerprinting diagnostic compounds, (e.g., saturated hydrocarbons, polynuclear aromatic hydrocarbons (PAHs), alkylated PAHs) by Alpha Analytical using United States Environmental Protection Agency (EPA) method 8015 (GC-FID; saturates) and a modified method 8270D (GCMS; PAHs). Full Methods details are provided in Stout 2016.

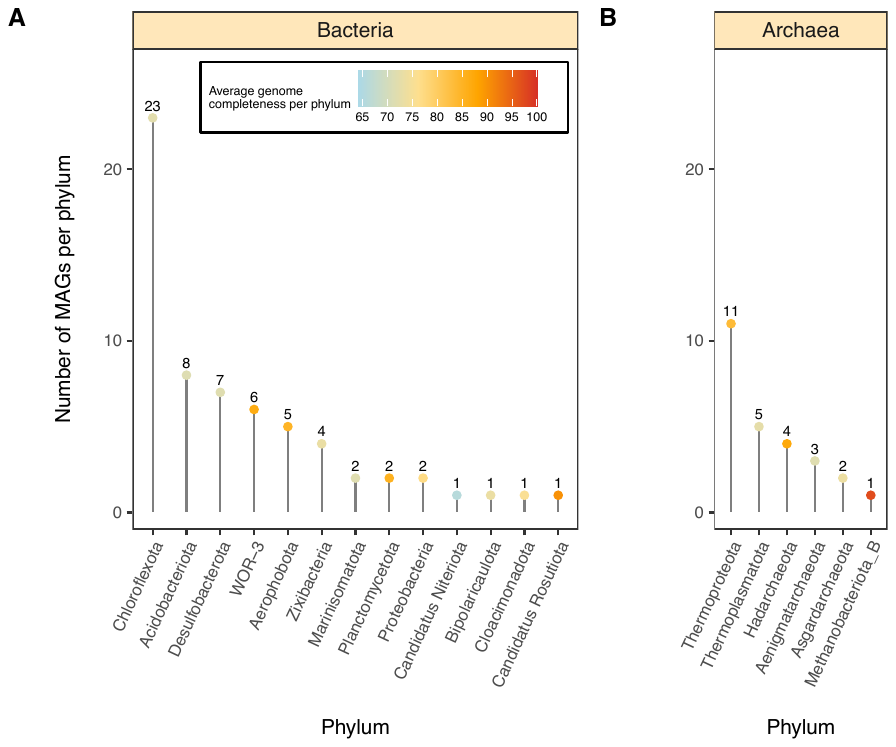

**Supplementary Figure 2**. **MAG recovery frequency**. Frequency of Guaymas Basin prokaryotic MAGs (≥ 50% completeness, ≤ 10% contamination) by bacterial (**A**) and archaeal phylum (**B**). Colored dots at the end of each line segment correspond to the mean genome completeness of the phylum; the number above the dot quantifies the number of genomes recovered from the phylum.

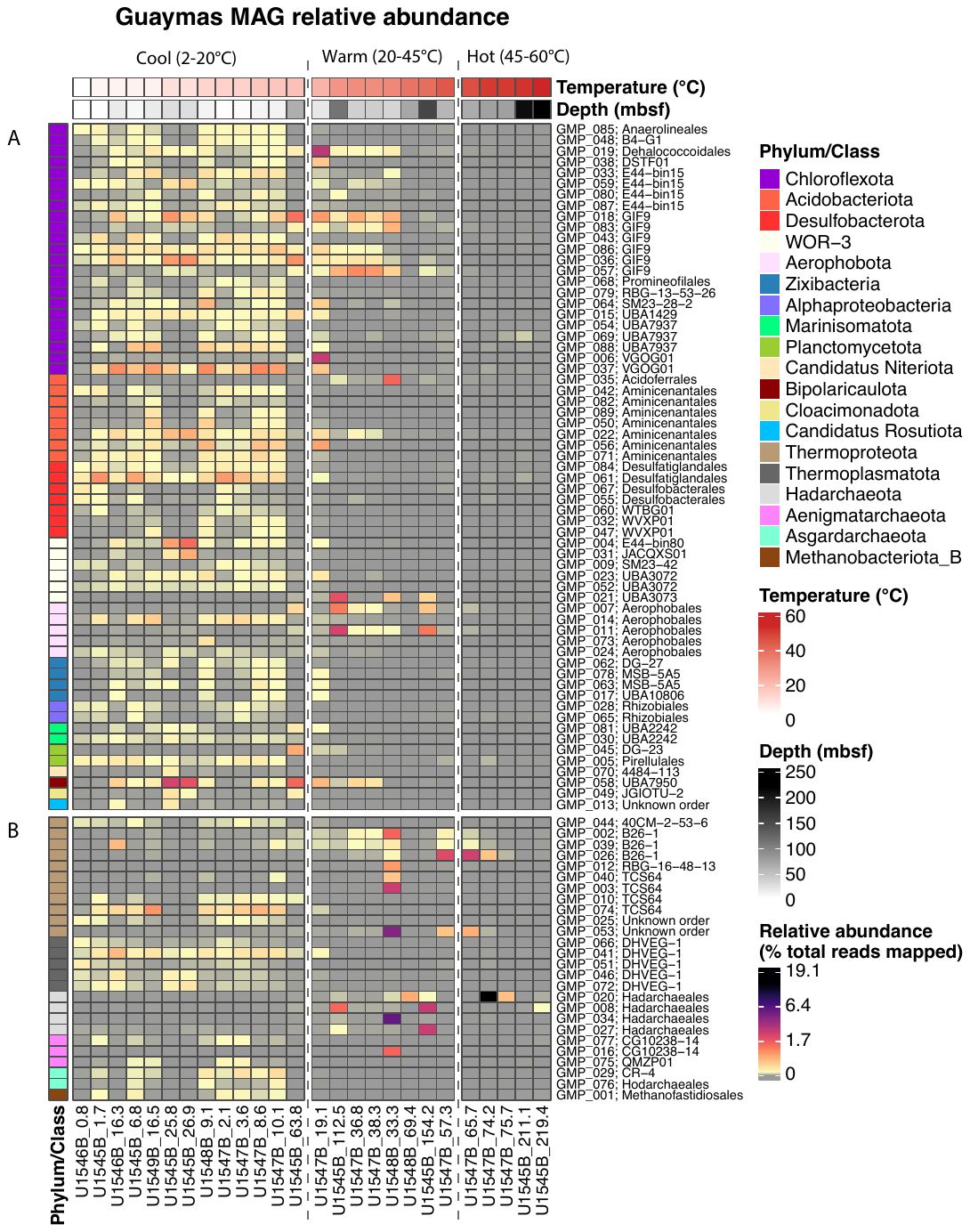

**Supplementary Figure 3. MAG relative abundances across taxonomic orders, modified from Figure 2.** The heatmap shows for each column the percentage of total pre-processed reads from a metagenomic sample that mapped to all 89 MAGs (across all samples) in order of increasing temperature from left to right. Temperature regimes (Cool, Warm, and Hot) are separated by vertical dashed lines. The abbreviated MAG names and their taxonomic orders are listed to the left of each row. “GMP_” is an abbreviation for the prokaryotic Guaymas MAG naming scheme “Guaymas_MAG_P_”. Each row represents the abundance profile of an individual MAG across all samples. MAGs are color coded using a colored annotation column to the left of the plot; taxonomic phylum/class names are given in the legend to the right of the heatmap. Samples are color coded using a colored annotation row that displays increasing temperature from left to right, and the associated depth (mbsf) each sample was collected from. The x-axis labels indicate sites and depths in mbsf.

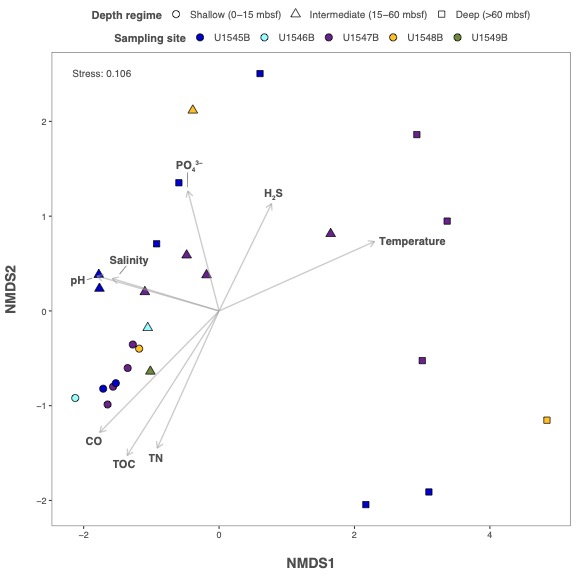

**Supplementary Figure 4. NMDS analysis of metagenomes and environmental parameters.** Non-metric multidimensional scaling (nMDS) ordination plot of metagenomes reconstructed from sites U1545B, U1546B, U1547B, U1548B, U1549B and environmental parameters with a significant effect on microbial community composition. The nMDS plot depicts the separation of Guaymas Basin metagenomic samples drilled from depths ranges for shallow (0-15 mbsf), intermediate (15-60 mbsf) and deep (> 60 mbsf) sediments, and statistically significant (p < 0.05) environmental parameters; plot stress: 0.106. The directions of the arrows indicate a positive or negative correlation among the environmental parameters with the ordination axes. The length of the arrow reflects the strength of correlation between the environmental parameter and the microbial community composition of metagenomic samples, with longer lines indicating stronger correlations.

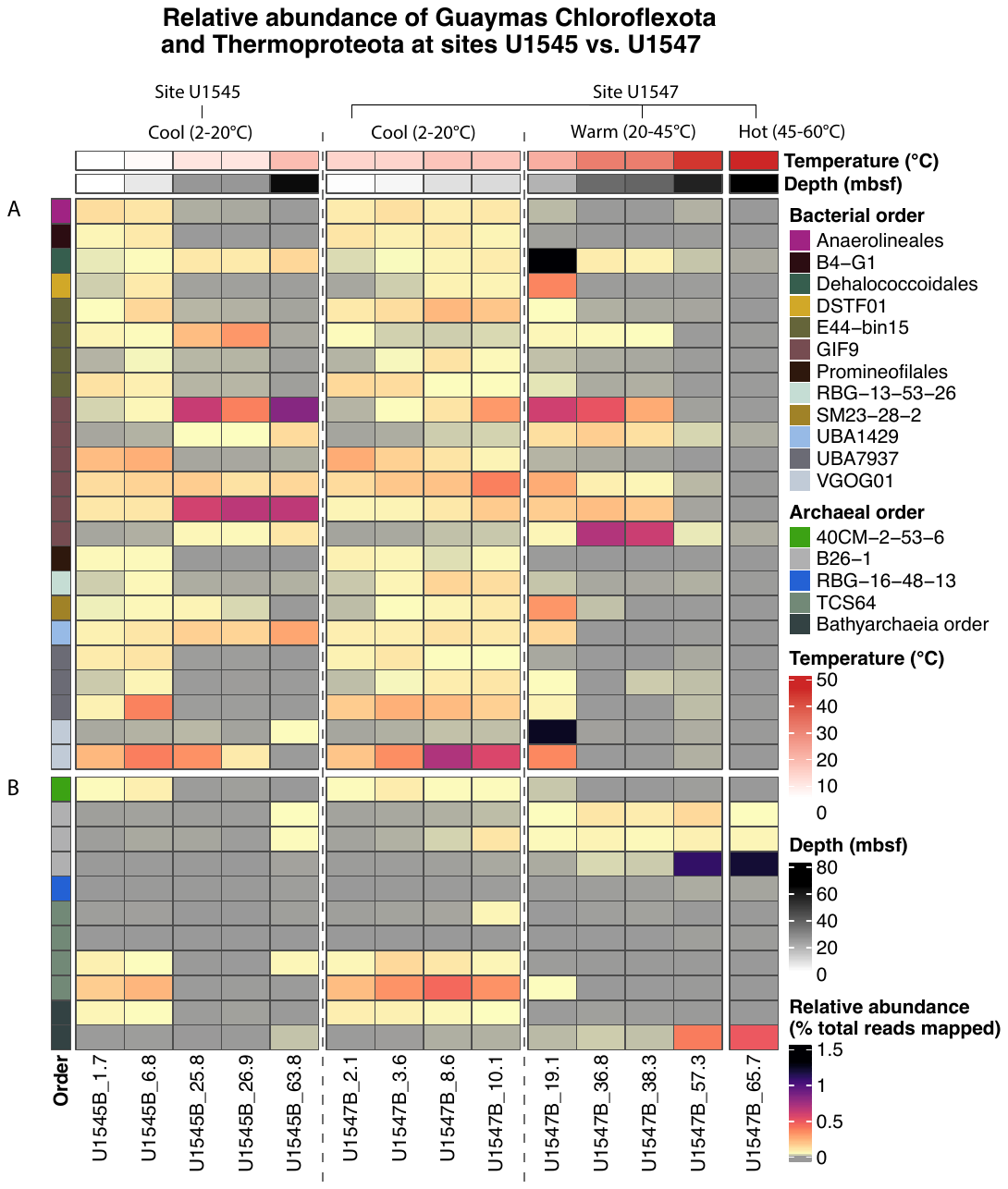

**Supplementary Figure 5. Relative abundance of Chloroflexota and Thermoproteota MAGs, identified by order, in metagenomic samples from drill sites U1545B and U1547B.** The heatmap for Chloroflexota (**A**) shows for each column the percentage of total pre-processed reads from a metagenomic sample that mapped to all Chloroflexota MAGs in order of increasing sampling depth from left to right for samples from sites U1545B (left) and U1547B (right). We show only depths that we had samples from both sites (to facilitate abundance pattern comparisons between sites). In the heatmap for Thermoproteota MAGs (**B**), the first 5 columns from the right-hand side correspond to site U1545B samples; followed by the site U1547B samples in the remaining columns after the gap between columns 5 and 6. MAG taxonomy is color coded to the left of the plot; taxonomic order names are given in the legend to the right of the heatmap. Samples are ordered by increasing temperature (left to right) at both sites, and temperatures and depths are color coded at the top of the plot.

**Table legends**

**Supplementary Table 1. Geochemical data from sites and depths where metagenomic data are available.** The columns from left to right include: sample (the name of the metagenomic sample), Depth (mbsf), Temperature, pH, Alkalinity (mM), Salinity, SO_4_^2-^ (mM), PO_4_^3-^ (μM), H_2_S (μM), NH_4_^+^ (mM), Mg^2+^ (mM), Ba^2+^ (μM), CH_4_ (mM), H_2_ (nM), CO (nM), DOC (mM), DIC (mM), CaCO_3_ (wt%), TOC (wt%), TN (wt%), TOC/TN, Fe (μM).

**Supplementary Table 2. Taxonomy and assembly statistics of removed putative contaminant MAGs.** Column 1 "bin_name" indicates the name of the contaminant bin, column 2 "checkm2_completeness" denotes the estimated completeness of MAGs using CheckM2, column 3 "checkm2_contamination" gives the estimated MAG contamination using CheckM2 (Chklovski et al., 2022a), column 4-10 give the taxonomic classifications for the MAGs: "domain", "phylum", "class", "order", "family", "genus", "species". Column 11 "das_tool_bin_score" denotes the Das Tool bin quality score for each MAG, column 12 "das_tool_scg_completeness" denotes the Das Tool bin completeness based on presence/absence of single copy genes (genes occurring only once in a specific taxonomic group/domain) within each MAG, column 13 "das_tool_scg_redundancy" denotes the number of Das Tool (Sieber et al., 2018) single copy gene duplicates in each MAG, column 14 "genome_size" gives the genome size in base pairs of each MAG, column 15 "n_contigs" gives the number of contigs in each MAG, and column 16 "N50" gives the N50 score of each MAG.

**Supplementary Table 3. Taxonomy and assembly statistics of all 89 metagenome-assembled genomes**. Column 1 "mag_name" indicates the name of the MAG, column 2 "checkm2_completeness" denotes the estimated completeness of MAGs using CheckM2, and column 3 "checkm2_contamination" gives the estimated MAG contamination using CheckM2 (Chklovski et al., 2022a). Column 4-10 give the taxonomic classifications for the MAGs: "domain", "phylum", "class", "order", "family", "genus", "species". Column 11 "das_tool_bin_score" denotes the Das Tool (Sieber et al., 2018) bin quality score for each MAG, column 12 "das_tool_scg_completeness" denotes the Das Tool bin completeness based on presence/absence of single copy genes (genes occurring only once in a specific taxonomic group/domain) within each MAG, column 13 "das_tool_scg_redundancy" denotes the number of Das Tool (Sieber et al., 2018) single copy gene duplicates in each MAG; column 14 "genome_size" gives the genome size in base pairs of each MAG, column 15 "n_contigs" gives the number of contigs in each MAG, and column 16 "N50" gives the N50 score of each MAG.

**Supplementary Table 4. KofamScan gene annotations for all medium quality MAGs (≥ 50% completeness, ≤ 10% contamination: *n* = 89).** Column 1 “mag_name” contains the name of all bacterial and archaeal MAGs (*n* = 89) from this study; columns 2-8 list the taxonomic domain, phylum, class, order, family, genus, and species of that the MAG containing the genome annotation belongs to; column 9 “adaptive_threshold” denotes whether an HMM score surpassed the adaptive scoring threshold or not with an asterisk (*); column 10 “gene_name” contains the name of the predicted gene from the output of Prodigal gene calling on each of the MAGs (the gene header names from each of the output amino acid fasta files containing predicted genes). Column 11 “k_number” contains the KEGG Orthology K number for each gene annotation; column 12 “e_value” denotes the E-value of the HMM alignment; column 13 “score” contains the HMM alignment score (the log odds ratio of the alignment to a particular protein family relative to a random model); column 14 “threshold” denotes the suggested HMM alignment score threshold for a robust annotation; column 15 “KO definition” contains the full gene name information for the annotation. Column 16 “score_type” indicates whether the HMM models the full protein sequence or a specific protein domain.

**Supplementary Table 5. Tab a: Prokka gene annotations for all medium quality MAGs (≥ 50% completeness, ≤ 10% contamination: *n* = 89).** Column 1 “mag_name” contains the name of all bacterial and archaeal MAGs (*n* = 89) from this study; column 2 “gene_name” contains the name of the predicted gene from the output of Prodigal gene calling on each of the MAGs (the gene header names from each of the output nucleotide fasta files containing predicted genes); columns 3-4 “start” and “end” contain the start and end positions of each gene sequence; column 5 “strand” indicates which strand of DNA the gene was on; columns 6-7 contain the gene name abbreviations (if any) and full gene name descriptions (“hypothetical protein” denotes an unknown protein); column 8 “dbxref” denotes which database was used to make the gene annotation; column 9 “ec_number” is the enzyme commission number; column 10 “contig” denotes the name of the contig the gene is located on; columns 11-17 list the taxonomic domain, phylum, class, order, family, genus, and species affiliations (if any) of the corresponding MAG. **Tab b:** **METABOLIC gene annotations for all medium quality MAGs (≥ 50% completeness, ≤ 10% contamination: *n* = 89).** Column 1 “mag_name” contains the name of all bacterial and archaeal MAGs (*n* = 89) from this study; columns 2-3 contain the taxonomic phylum and order of each of the 89 MAGs; column 4 “Category” contains the metabolism category associated with the gene annotation; column 5 “Function” indicates which enzymatic functional group the gene annotation is associated with; columns 6-7 contain the gene name abbreviations and full gene name descriptions; column 8 “Hmm.file” denotes which HMM files were used to make the gene annotation; column 9 “Corresponding.KO” is the KEGG ortholog identifier; column 10 “Reaction” denotes the reaction the encoded enzyme is involved in; column 11 “Substrate” indicates involved substrates in the reaction; column 12 lists the product(s) of the reaction; column 13 “Hmm.detecting.threshold” denotes the threshold cutoffs for each hmm; column 14 "Hits” denotes the gene that received a hit to the HMMs .

**Supplementary Table 6. AntiSMASH 6.0 biosynthetic gene cluster annotations for all medium quality MAGs (≥ 50% completeness, ≤ 10% contamination: *n* = 89).** Column 1 “mag_name” contains the name of all bacterial and archaeal MAGs that contained BGCs > 5 kilobases in length; columns 2-5 contain the taxonomic domain, phylum, class, and order of each of the MAGs; column 6 “bgc_name” contains the biosynthetic gene cluster name; columns 7-8 contain the start and end locations of each of the genes in the biosynthetic gene cluster; column 9 “strand” indicates which strand of DNA the gene was on; column 10 “product” contains the biosynthetic product encoded by the cluster; column 11 “length” contains the length of the gene cluster in base pairs, column 12 “contig” indicates which contig the cluster was detected on.

**Supplementary Table 7**. **Output table of MetaPathPredict depicting the predicted presence or absence of KEGG metabolic modules (*n* = 476) for Cloacimonadota and Zixibacteria bacterial MAGs (*n* = 5).** Column 1 “module_class” indicates the category the KEGG module belongs to (if any, otherwise it is labelled as “No module class”); column 2 “module_name” gives the name of each KEGG module; column 3 “module_number” gives the module numeric identifier used in the KEGG database; each of the remaining 5 columns contain all 476 KEGG module presence/absence predictions for each Cloacimonadota and Zixibacteria MAG. The follow classifications are used in the color-coded cells of these columns: “present: predicted” (the module is present in the genome and was predicted present by MetaPathPredict), “present: not predicted” (the module is present in the genome and was predicted absent), “absent: not predicted” (the module is completely absent in the genome and was predicted absent), “absent: predicted” (the module is completely absent in the genome and was predicted present), “incomplete: not predicted” (the module is partially present in the genome and was predicted absent), and “incomplete: predicted” (the module is partially present in the genome and was predicted present).
